# Putative protective neural mechanisms in pre-readers with a family history of dyslexia who subsequently develop typical reading skills

**DOI:** 10.1101/707786

**Authors:** Xi Yu, Jennifer Zuk, Meaghan V. Perdue, Ola Ozernov-Palchik, Talia Raney, Sara D. Beach, Elizabeth S. Norton, Yangming Ou, John D. E. Gabrieli, Nadine Gaab

**Author notes:** **Corresponding author:** Xi Yu, PhD, Laboratories of Cognitive Neuroscience, Department of Medicine/Division of Developmental Medicine, Boston Children’s Hospital/Harvard Medical School, 1 Autumn Street (Office 641), Boston, MA 02115, USA.

## Abstract

Developmental dyslexia is a learning disability characterized by difficulties in word reading. While the prevalence in the general public is around 10-12%, an increased prevalence of 40-60% has been reported for children with a familial risk. Neural atypicalities in the reading network have been observed in children with (FHD+) compared to without (FHD-) a family history of dyslexia, even before reading onset. Despite the hereditary risk, about half of FHD+ children develop typical reading abilities (FHD+Typical) but the underlying neural characteristics and the developmental trajectories of these favorable reading outcomes remain unknown. Utilizing a retrospective, longitudinal approach, this is the first study to examine whether potential protective neural mechanisms are present before reading onset in FHD+Typical. Functional and structural brain characteristics were examined in 69 pre-readers who subsequently developed typical reading abilities (35 FHD+Typical/34 FHD-Typical) using MRI/fMRI. Searchlight-based multivariate pattern analyses identified distinct activation patterns during phonological processing between FHD+Typical and FHD-Typical in right inferior frontal (RIFG) and left temporo-parietal (LTPC) regions. Hypoactivation in LTPC was further demonstrated in FHD+Typical compared to FHD-Typical, suggesting that this previously reported neural characteristic of dyslexia is primarily associated with familial risk. Importantly, FHD+Typical pre-readers exhibited higher activation in RIFG than FHD-Typical, which was associated with increased interhemispheric functional and structural connectivity. These results suggest that putative protective neural mechanisms are already established in FHD+Typical pre-readers and may therefore support their successful reading development. Further studies are needed to investigate the functional significance and developmental trajectories of these neural mechanisms as well as their enabling factors, which has the potential to inform the design of early preventative/remediation strategies.

## Introduction

Developmental dyslexia (dyslexia) is a neurodevelopmental learning disability, which is characterized by difficulties with speed and accuracy of single word reading, deficient decoding abilities and poor spelling (IDA, 2007). Dyslexia has further been associated with functional and structural alterations in primarily left-hemispheric reading network components comprising the frontal, temporoparietal and occipito-temporal areas that underlie typical reading abilities (McCandliss & Noble, 2003; Norton et al., 2015; Ozernov-Palchik et al., 2016; Pugh et al., 2000; Richlan et al., 2009, 2011, 2013). Children with dyslexia often experience severe difficulties in their academic and personal lives as well as mental health due to the importance of reading in school curricula and in society (Baker & Ireland, 2007; Dougherty, 2003; Morgan et al., 2008).

There is an increased familial occurrence of dyslexia, ranging from 40% to 60%, compared to a prevalence of around 10% in the general population (Astrom et al., 2007; Katusic et al., 2001; Snowling & Melby-Lervåg, 2016) and several susceptibility genes have been identified (e.g., Newbury et al., 2011; Poelmans et al., 2011; Scerri et al., 2011; Taipale et al., 2003). Behavioral longitudinal studies have demonstrated early deficits in language and preliteracy skills (e.g., phonological processing and rapid automatized naming) in toddlers and preschoolers with (FHD+) compared to without (FHD-) a familial risk of dyslexia (e.g., Koster et al., 2005; Lyytinen et al., 2004; Lyytinen et al., 2001; Plakas et al., 2013; van der Leij et al., 2013). Moreover, neural alterations at the functional and structural levels have also been observed in FHD+ children before reading onset, and as early as in infancy (Guttorm et al., 2001; Im et al., 2015; Langer et al., 2017; Leppänen et al., 1999; Maurer et al., 2003; Raschle et al., 2011, 2012, 2013; van Herten et al., 2008; van Leeuwen et al., 2006; Vandermosten et al., 2015; Wang et al., 2016), also see (Ozernov-Palchik & Gaab, 2016) for a review). The early emergence of cognitive and neural alterations in FHD+ children suggests that the observed atypicalities might serve as developmental mechanisms that contribute to dyslexia susceptibility instead of being the result of reduced reading experiences (e.g., Lyytinen et al., 2001; Raschle et al., 2012; Snowling & Melby-Lervåg, 2016). Nevertheless, approximately half of FHD+ children subsequently develop typical reading skills (Gallagher et al., 2000; Pennington & Lefly, 2001; Scarborough, 1990; Snowling et al., 2003; Snowling & Melby-Lervåg, 2016; Torppa et al., 2010). Previously, longitudinal behavioral studies have identified several protective factors in FHD+ pre-readers that support their subsequent typical reading development, including enhanced oral language abilities, particularly in vocabulary knowledge and syntactic structure, and increased executive functioning skills (e.g., Haft et al., 2016; Plakas et al., 2013; Snowling et al., 2003; Snowling & Melby-Lervåg, 2016; Torppa et al., 2010). However, it remains unknown whether there are also protective mechanisms in the pre-reading brain that may facilitate typical reading development in FHD+Typical children.

Compensatory neural mechanisms have previously been investigated in older struggling readers after several years of formal reading instruction. These studies, in general, suggest that the learning difficulties in reading experienced by these children might be mediated by neural compensatory pathways involving the right-hemispheric (RH) network (Barquero et al., 2014). More specifically, increased activation in RH regions have been shown in compensated readers compared to those with persistently poor reading skills (Shaywitz et al., 2003) and in individuals who demonstrated reading improvement after successful interventions (e.g., Eden et al., 2004; Temple et al., 2003). Moreover, right-hemispheric neural characteristics of struggling readers, such as increased neural activation in the right frontal cortex during phonological processing and stronger connectivity strength of the right white matter tracts important for reading, have also been shown to predict their subsequent reading improvement (Farris et al., 2016; Hoeft et al., 2011). In addition, the compensatory role of RH has further been observed in children with dyslexia, for which higher neural sensitivities for speech sounds have been associated with better phonological and reading skills (Lohvansuu et al., 2014).

In addition to the development of compensatory mechanisms in poor readers or children with dyslexia, which are probably developing in response to successful reading intervention, it has been hypothesized that some children might already exhibit protective neural mechanisms in the right hemisphere at the pre-reading stage. This may enable them to develop typical reading skills that might be otherwise compromised due to atypical/alternative brain development associated with a familial risk for dyslexia (Yu et al., 2018). As a group, infants and children with a familial risk of dyslexia seem to show a greater predisposition for a bilateral/right-lateralized brain network involved in language and reading development, compared to controls who show primarily a left-hemispheric dominance (e.g., Guttorm et al., 2001; Leppänen et al., 1999; Lyytinen et al., 2005; van Herten et al., 2008; van Leeuwen et al., 2006; van Leeuwen et al., 2007; Wang et al., 2016; for reviews, also see Lyytinen et al., 2005 and Ozernov-Palchik & Gaab, 2016). For example, enhanced neural sensitivity to speech sounds in the right hemisphere has been observed in FHD+ compared to FHD-infants within the first couple days of life (Guttorm et al., 2001). In a recent longitudinal study examining white matter development from the prereading (before kindergarten entry) to the fluent reading stage (up to 5^th^ grade), right lateralization in white matter tracts important for reading has also been observed in FHD+ compared to FHD-preschoolers (Wang et al., 2016). Importantly, faster white matter development in the right hemisphere has been demonstrated in subsequent good versus poor readers within a group of FHD+ children, suggesting possible early neural compensatory mechanisms in the right hemisphere. However, it remains unclear whether these alternative RH neural pathways are only developed to compensate for the difficulties/impairment children encounter after they start to learn to read (i.e., **compensatory mechanisms**), or whether they are already in place prior to reading onset (e.g., at birth or in early childhood) in some FHD+ children and thereby provide a protective role from the start of learning to read (i.e., **protective mechanisms**, Yu et al., 2018).

Alternatively, one could also argue that typical reading development among FHD+Typical children may be simply the result of lower genetic liability compared to children who have a familial risk and exhibit reading impairment (FHD+Impaired, Snowling et al., 2003; Van Bergen et al., 2011). Behavioral studies tracking FHD+ and FHD-children longitudinally over the course of learning to read have indicated that the liability for dyslexia is a continuous variable among children with familial risk. Specifically, FHD+Impaired children have shown lower performance on the key precursors of dyslexia, including phonological awareness, automatized rapid naming skills and letter knowledge, compared to FHD+Typical children. However, FHD+Typical children have also been shown to perform more poorly than FHD-Typical children on these pre-literacy and reading measures (Pennington & Lefly, 2001; Snowling & Melby-Lervåg, 2016), indicating that the liability for dyslexia is a continuum and not a dichotomic variable. Nevertheless, it is unknown whether (a) FHD+Typical children display the characteristic functional and structural brain alterations previously described for children with a diagnosis of dyslexia due to their genetic liability, (b) what are the neural protective/compensatory mechanisms associated with typical reading development and (c) are these potential mechanisms present prior to reading onset which would suggest that they play a protective role.

Utilizing a retrospective, longitudinal approach, the current study is the first to investigate whether putative protective neural mechanisms emerge prior to reading onset in preschoolers and early kindergarteners with a familial risk of dyslexia who subsequently develop typical reading skills. Sixty-nine typically reading children (35 FHD+Typical and 34 FHD-Typical) were selected from our longitudinal database based on their reading performance, which had been assessed after having received at least two years of reading instruction. Retrospective analyses of the behavioral, structural (diffusion), and functional (phonological processing) data collected at the pre-reading stage (before or at the beginning of their kindergarten) were conducted. FMRI data were first subjected to a mass-univariate analysis as well as a searchlight multivariate pattern analysis (MVPA) to explore potential group differences in individual voxels and activation patterns across neighboring voxels, respectively. Regions showing distinct activation patterns between the FHD-Typical and FHD+Typical children, as identified by the MVPA, were further characterized for the specific pattern associated with each group. Functional connectivity (FC) analyses were subsequently performed to investigate network characteristics in FHD+Typical compared to FHD-Typical children. Additionally, in order to identify potential protective mechanisms in the white matter microstructure, FA values in right-hemispheric white matter tracts previously associated with reading skills (Horowitz-Kraus et al., 2014), especially in compensated readers (Hoeft et al., 2011; Wang et al., 2016) were compared between groups (via two-sample t-tests); these tracts included the right arcuate fasciculus (RAF), inferior longitudinal fasciculus (RILF) and superior longitudinal fasciculus (RSLF). The corpus callosum (CC) was also examined since it plays a critical role in interhemispheric communication and brain lateralization (Aboitiz & Montiel, 2003; Hinkley et al., 2016) and whiter matter microstructure of the CC has been associated with variants of dyslexia susceptibility genes (Darki et al., 2012; Scerri et al., 2012).

Overall, we hypothesize that if FHD+Typical children develop typical reading skills as a result of a reduced/null liability, we will not observe different brain mechanisms underlying reading development between FHD- and FHD+ children who subsequently developed equivalent, typical reading abilities. Alternatively, one can hypothesize that FHD+Typical children may exhibit neural deficits in the left-hemispheric reading network as a result of their familial risk, but may utilize protective pathways and mechanisms in RH brain regions through increased interhemispheric functional and structural (CC) connectivity, which may already be established prior to reading onset to support their successful reading development.

## Methods

### General study design

The current study was based on two longitudinal projects in our lab; the ‘Boston Longitudinal Dyslexia Study’ (BOLD) and ‘Research on the Early Attributes of Dyslexia’ (READ) at Boston Children’s Hospital (BOLD and READ) and the Massachusetts Institute of Technology (READ). In both projects, children were initially enrolled at the end of the pre-kindergarten or the beginning of kindergarten before they started to learn to read (i.e., pre-readers), where they completed both behavioral and imaging sessions (more details provided below). All participants were then longitudinally followed to track reading development until the end of second grade (READ) or fourth grade (BOLD).

### Participants

An initial group of 93 participants, including 52 FHD+ children, defined as having at least one first-degree relative with a dyslexia diagnosis, and 41 FHD-controls were retrospectively selected from both longitudinal projects using the following criteria: 1) neural and behavioral data successfully collected at the pre-reading stage (see below for details); 2) reading skills subsequently assessed at the emergent reading stage. Among these children, 12 FHD+ (24%) and 2 FHD-(5%) children showed reading disabilities, since they scored lower than the clinical cutoff (standard scores (SS) < 85) in any of the four word-level reading assessments, including the Word ID (WI) and Word Attack (WA) subtests of the Woodcock Reading Mastery Test-Revised (WRMT-R), Woodcock, 1987), as well as the Sight Word Efficiency (SWE) and Phonemic Decoding Efficiency (PDE) subtests of the Test of Word Reading Efficiency (TOWRE, Torgesen et al., 1999) during their latest assessment session. The prevalence of dyslexia was higher in FHD+ compared to FHD-children (χ2(1) = 5.94, p = 0.015), which was consistent with previous literature (e.g., Boets et al., 2007; Torppa et al., 2010; Van Bergen et al., 2012). To ensure that participants in the current study were pre-readers at the initial stage, 10 more children (5 FHD-Typical and 5 FHD+Typical) who identified more than 10 words (25.1 words ± 11.8, range = 11-43) on the WI subtest of the WRMT-R were further excluded. The final sample included 34 (18 males) FHD+Typical and 35 (18 males) children. All participants were native English speakers and exhibited non-verbal IQs of SS > 80, as measured by the Kaufman Brief Intelligence Test: Second Edition – Matrices (KBIT-2), Kaufman & Kaufman, 2004). Most children were right handed, with three left-handed children (1 FHD+Typical and 2 FHD-Typical) and one child (FHD-Typical) who did not demonstrate a preference (ambidextrous) was also included in the sample. No children reported a history of hearing, vision, psychiatric or neurological disorders. The current study was approved by the institutional review boards at the Massachusetts Institute of Technology and Boston Children’s Hospital. Before participation, verbal assent and informed written consent were obtained from each child and guardian, respectively.

## Longitudinal psychometric measurements

At the pre-reading stage, all children were assessed on pre-literacy skills including (a) phonological processing (the Comprehensive Test of Phonological Processing (CTOPP, Wagner et al., 1999), which measures the ability to segment, combine and repeat phonological components, (b) rapid automatized naming abilities (RAN, Wolf & Denckla, 2005), which indicates automaticity of phonological access through measuring the amount of time it takes a child to name a series of symbols (e.g., objects and numbers) as fast as possible, and (c) letter knowledge (the Letter Identification subtest of WRMT-R). Their non-verbal IQ (KBIT-2) and word reading abilities (WRMT-R, Word ID) were also evaluated. Moreover, children from the BOLD project were further examined on their language skills using the Clinical Evaluation of Language Fundamentals: Fourth edition (CELF-4, Semel et al., 2003). Home literacy environment (adapted from Denney et al., 2001), supplementary Table S1) and socioeconomic status (adapted from the MacArthur Research Network: http://www.macses.ucsf.edu/default.php, supplementary Table S2) were characterized based on parent reporting at the first study visit. Children’s reading abilities were assessed at the emergent reading stage using the WI and WA subtests of the WRMT-R, as well as the SWE and PDE subtests of the TOWRE. Although the number of acquired time points varied among participants due to scheduling challenges in longitudinal studies, all children included in the current study were successfully assessed at least once after two years of formal reading instruction (i.e., the end of 1^st^ grade). The latest available reading performance for each child (ranging from the end of 1^st^ and 4^th^ grade) was used for the current analyses. The FHD-Typical and FHD+Typical groups did not differ in age and school grade associated with the latest available reading performances (grade: *χ*2(3) = 5.50, *p* = 0.14; age: *t*_67_ = 1.2, *p* = 0.23, see information on grade distribution and age for each group in Table 1).

**Table 1.** Pre-literacy characteristics, fMRI experiment performance conducted at the pre-reading stage, and reading abilities after schooling.

|  | FHD-Typical | FHD+Typical | Group Effect |
| --- | --- | --- | --- |
| Number (female/male) | 34 (16/18) | 35 (17/18) | - |
| **Pre-literacy characteristics** |  |  |  |
| Age (months) | 65 ± 4.3 | 66 ± 4.7 | $t_{67} = 1.3; p = 0.2$ |
| CTOPP: Elision | 10 ± 2.1 | 10 ± 2.4 | $t_{64} = 0; p = 1$ |
| CTOPP: Blending | 11 ± 1.9 | 11 ± 2.2 | $t_{64} = 0.3; p = 0.8$ |
| CTOPP: Nonword Repetition | 9.4 ± 1.6 | 9.3 ± 2.0 | $t_{65} = 0.3; p = 0.8$ |
| RAN: Object | 104 ± 10 | 100 ± 13 | $t_{62} = 1.5; p = 0.1$ |
| RAN: Color | 101 ± 13 | 96 ± 17 | $t_{62} = 1.2; p = 0.2$ |
| WRMT-R: Word ID | 93 ± 15 | 96 ± 22 | $t_{67} = 0.6; p = 0.5$ |
| KBIT-2: non-verbal | 103 ± 11 | 99 ± 9.8 | $t_{67} = 1.6; p = 0.1$ |
| CELF-4: Core Language | 113 ± 14 | 110 ± 10 | $t_{58} = 1.0; p = 0.3$ |
| CELF-4: Receptive Language | 111 ± 13 | 104 ± 15 | $t_{59} = 1.7; p = 0.1$ |
| CELF-4: Expressive Language | 114 ± 14 | 110 ± 12 | $t_{58} = 1.2; p = 0.2$ |
| CELF-4: Language Structure | 114 ± 15 | 110 ± 11 | $t_{57} = 1.4; p = 0.2$ |
| **Reading abilities at the end of the first grade or later** |  |  |  |
| # of participants with latest reading performance available in each grade |  |  |  |
| First grade | 8 | 5 | $\chi^2_{(2)} = 1.5$ |
| Second grade | 17 | 17 |  |
| Third grade | 1 | 7 |  |
| Fourth grade | 8 | 6 |  |
| Age (months) | 104 ± 14 | 108 ± 13 | $t_{67} = 1.2; p = 0.2$ |
| WRMT-R: Word ID | 111 ± 9.7 | 108 ± 10 | $t_{65} = 1.2; p = 0.2$ |
| WRMT-R: Word Attack | 109 ± 11 | 109 ± 10 | $t_{65} = 0.05; p = 0.95$ |
| TOWRE: SWE | 109 ± 13 | 104 ± 9.8 | $t_{67} = 1.7; p = 0.1$ |
| TOWRE: PDE | 104 ± 11 | 104 ± 9.6 | $t_{67} = 0.02; p = 0.98$ |

Note. Standard scores were reported for all the psychometric assessments. Due to the miss of data points in each assessment, degree of freedom and significance level were adjusted accordingly. No significant differences were observed between the FHD-Typical and FHD+Typical children for any of the measures described in the table.
CTOPP: Comprehensive Test of Phonological Processing; RAN: Rapid Automatized Naming; KBIT-2: Kaufman Brief Intelligence Test: Second Edition--nonverbal matrices; CELF: Clinical Evaluation of Language Fundamentals: Fourth Edition; WRMT-R: Woodcock Reading Mastery Tests-Revised; TOWRE: Test of Word Reading Efficiency; SWE: Sight Word Efficiency; PDE: Phonemic Decoding Efficiency.

Raw scores initially acquired from each assessment were converted into standard scores (Mean (M): 100, Standard Deviation (SD): 15; scale scores (M: 10, SD: 3) for the CTOPP results) for result summaries and statistical analyses. To evaluate any potential difference in behavioral performance between the FHD-Typical and FHD+Typical groups, a two-sample t-test was carried out for each of the psychometric measurements collected at the pre-reading and emerged reading stages. Similarly, chi-squared tests were performed to evaluate group differences in home literacy environment and socioeconomic status. Significant group differences were reported at *p* < 0.05,

## Imaging experiment at the pre-reading stage

### Imaging acquisition

Neuroimaging data collection for the BOLD and READ projects were conducted at Boston Children’s Hospital (BCH) and Massachusetts Institute of Technology (MIT), respectively. Images were acquired on a 3T Siemens Trio Tim MRI scanner with a standard Siemens 32-channel phased array head coil at both sites. For fMRI data collection, a behavioral interleaved gradient imaging design was applied to minimize the influence of scanning background noise during auditory stimulus presentation (Gaab et al., 2007a, 2007b), using the following parameters: TR = 6,000 ms; TA = 1,995 ms; TE = 30 ms; flip angle = 90°; field-of-view = 256 mm^2^; in-plane resolution = 3.125×3.125 mm^2^, slice thickness = 4 mm, slice gap = 0.8 mm). Structural images were acquired using site-specific specification as follows: for BCH, slice number = 128, TR = 2000 ms, TE = 3.39 ms, flip angle = 9°, field of view = 256 mm^2^, voxel size = 1.3×1.0×1.3 mm^3^; for MIT, slice number = 176, TR= 2350 ms, TE= 1.64 ms, flip angle=9°, FOV= 256 mm^2^, voxel size 1.0 x 1.0 x 1.0 mm^3^. Finally, DTI scans collected at both sites included 10 nondiffusion-weighted volumes (b=0) and 30 diffusion-weighted volumes acquired with non-colinear gradient directions (b=1000 s/mm^2^ for BCH and b=700 s/mm^2^ for MIT), all at 128×128 base resolution and isotropic voxel resolution of 2.0 mm^3^. Note that only half of the BOLD participants have DTI data available since this sequence was only added later during the BOLD recruitment process. Throughout the MRI session there was one research assistant accompanying the child participant to ensure minimal head movement and compliance with the task instructions (see detailed protocol in Raschle et al., 2009, 2012).

### Task procedure

A phonological processing task was presented in block design using Presentation software (Version 0.70, www.neurobs.com). During each trial, children saw two common objects presented on the left and right sides of the screen sequentially (2 seconds for each), while hearing each object’s name, spoken in a male or female voice, accompanying their visual appearance. The two pictures stayed on the screen for an additional 2 seconds, while participants judged whether the first sound of the names of the two objects matched (first sound matching, FSM) by pressing buttons held in the right (Yes) and left (No) hands. The FSM run was comprised of seven task blocks, each consisting of four trials, alternating with seven fixation blocks of the same length (24 seconds). A separate experimental run with a control task was constructed with the same stimuli in the same way; however, in this run, participants were asked to decide whether the object names were spoken by the same gender (voice matching, VM). The order of the two runs was counterbalanced across participants (see more details in Raschle et al., 2012, 2013).

### In-scanner performance

Participants’ responses were recorded during the neuroimaging session. Given the young ages of the participants, self-correction was allowed, and only the last response within each trial was used to compute the number of correct responses and RTs. Group differences in the in-scanner performance were evaluated using two-sample t-tests.

### FMRI analyses

fMRI data were successfully collected in 30 FHD-Typical and 30 FHD+Typical children. Acquired images were first preprocessed in SPM8 (http://www.fil.ion.ucl.ac.uk/spm/software/spm8) based on Matlab (Mathworks) using an age-appropriate pipeline (Yu et al., 2018). Specifically, after removing the initial volumes due to the T1 equilibration effects, functional images were first motion corrected (realigned) and co-registered to the corresponding structural images. Before normalizing fMRI images from the naïve space to the Montreal Neurological Institute (MNI) space, transformational matrices were first estimated for every participant using their corresponding high-resolution structural images in VBM8 (http://www.neuro.uni-jena.de/vbm/). During this step, structural images were segmented into grey matter (GM), white matter (WM), and cerebral spinal fluid (CSF) using an adaptive Maximum A Posterior (MAP) approach (Rajapakse et al., 1997) and spatially normalized to the MNI space via affine transformation. An age- and gender-matched Tissue Probability Map created using the Template-O-Matic Toolbox (Wilke et al., 2008) was applied at this stage to accommodate the potential anatomical differences between the brain images of the current pediatric group and MNI templates created based on the adult population (Evans, 1992). A non-linear normalization step was subsequently performed on the GM and WM using a diffeomorphic anatomical registration using exponentiated lie algebra (DARTEL) approach (Ashburner, 2007). Another customized template created based on structural images of 149 children with a similar age (67.9 ± 4.2 months) and gender ratio (Female/Male = 1.04/1) to the current participant group were applied during DARTEL registration. The linear-transformed GM and WM were mapped to this template through high dimensional warping processes, resulting in optimal registration in local, fine-grained structures among all the participants. The transformational matrices from the native space to the MNI space were generated for each participant after VBM processing and then applied to the corresponding functional images for the normalization purpose. A Gaussian kernel with full-width at half maximum (FWHM) of 8mm was further applied to create smoothed images. Finally, to minimize the effect of head motion on data analyses, Artifact Detection Tools (http://www.nitrc.org/projects/artifact_detect) was applied to identify scans with excessive motion, using the criterion at 3mm (translational) and/or 2° (rotational). All the selected images were visually screened, and those with artifacts, such as missing voxels, stripes, ghosting, or intensity differences were marked as outliers and removed from subsequent analyses.

The preprocessed images were then entered into first-level general linear models for estimation of neural responses associated with task conditions (FSM, VM, i.e., regressors of interest). Comprehensive measurements of head motion along three translational and three rotational dimensions combined with the binary regressors representing outlier images were also included as covariates of no interest to minimize the confounding effect of head movement. The potential differences in the motion effect among the two groups were further evaluated through two-sample t-tests, which did not reveal any significant difference in the amount of absolute head movement in any of the six directions (Supplementary Table S4; 3 translational movement: left to right: *t*_58_ = 0.22, *p* = 0.83; posterior to anterior: *t*_58_ = 1.6, *p* = 0.11; bottom to top: *t*_58_ = 0.45, *p* = 0.66; 3 rotational movement: pitch: *t*_58_ = 0.34, *p* = 0.73; roll: *t*_58_ = 0.67, *p* = 0.51; yaw: *t*_58_ = 2.0, *p* = 0.06). Subject-wise neural responses for phonological processing were estimated by contrasting the beta map of the FSM condition with that of the VM condition for each participant. These contrast maps were subjected to the whole-brain group-level analyses to examine the differences in the functional mechanisms underlying phonological processing between FHD+Typical and FHD-Typical children.

#### Whole-brain analyses

were first performed to identify brain regions that showed activation differences between FHD+Typical and FHD-Typical groups. Two approaches were applied here. The first analysis utilized the mass-univariate analysis method to explore whether group differences in activation levels could be observed at the voxel level. To this aim, a two-sample t-test model was built and contrasts between the two groups were tested. In the second analysis, to capture group information embedded in the distributed patterns of brain activity, a searchlight MVPA (Kriegeskorte et al., 2006) was carried out using the TDT toolbox (Hebart et al., 2015). The MVPA is an analytic technique that surveys the relationships among the voxels (i.e., distributed representation) to identify activation patterns that differentiate experimental conditions (e.g., (Friston et al., 1994; Haxby et al., 2001; Norman et al., 2006). The combination of the MVPA with a searchlight technique further provided an opportunity for functional localization (Kriegeskorte et al., 2006). Specifically, a spherical searchlight was created for every voxel in the brain with a radius of 6 mm (2-voxel radius, resulting in 19 voxels in total). Then, for each searchlight, the contrast estimates were extracted from all included voxels for each participant and fed into a linear support vector classifier (SVC, LIBSVM—http://www.csie.ntu.edu.tw/~cjlin/libsvm). To make an unbiased estimation of the classification accuracies of the linear SVC, a 15-folder crossvalidation approach was adopted. All subjects were divided into 15 subgroups with two FHD+Typical and FHD-Typical children each. During each iteration, a classifier was trained on 14 subgroups (28 FHD-Typical and 28 FHD+Typical participants) and then used to predict the labels of the remaining two pairs. The process was repeated 15 times so that each subject was tested once, and the prediction accuracies of the SVC were estimated across all the subjects. Following Kriegeskorte et al. (2006) and Stelzer et al. (2013), the significance of the classification accuracies was determined by 5000 permutation tests in which group labels were randomly assigned to each subject for each searchlight. For both analyses, the statistical significances were further FDR-corrected for multiple comparisons, and region with a minimal of 5 connected voxels (mass-univariate) or searchlights (MVPA) showing *p*_corrected_ < .05 were reported. Moreover, to constrain the analyses to the cerebral cortex, a customized mask was created by overlapping a mean gray matter image (averaged across all the participants and thresholded at 0.1) with a cerebral mask derived from the Automated Anatomical Labeling atlas (Tzourio-Mazoyer et al., 2002).

Since the distinct activation patterns between groups could be reflected in the multi-voxel spatial pattern and/or a systemic difference across voxels (Jimura & Poldrack, 2012; Kragel et al., 2012), follow-up analyses were conducted in regions with significant MVPA results to evaluate whether the potential group differences in activation levels contributed to the distinct activation patterns observed between FHD-Typical and FHD+Typical children (Bauer & Just, 2017; Coutanche, 2013). To do this, the group differences in the contrast estimates for FSM > VM were computed for every voxel included in each searchlight. Meanwhile, the weight information of every feature (i.e., voxel) in the classification model was estimated for each significant searchlight using the whole data sets (30 FHD-Typical and 30 FHD+Typical). To account for the dependencies across the neighboring voxels, weight information was further corrected using the covariance matrix among all the voxels included in a searchlight (Haufe et al., 2014). The absolute value of the corrected weight, reflecting the true contribution magnitude of each voxel to the final classification performance, was then correlated with the group difference in activation levels across all the voxels in each searchlight. Statistical significance was held at *p*_corrected_ < 0.05, corrected for multiple comparisons.

Finally, the classification performance of each identified region as a whole (as compared to the individual searchlights it comprised) was evaluated in a ROI fashion. Specifically, all the connected searchlights from each significant region were combined into a cluster. The MVPA was performed following the same procedure as that in each searchlight, and the significance of the classification performance was evaluated using the permutation tests (n=5000). Furthermore, correlation analyses were also performed between the corrected weights of participating voxels and the corresponding voxel-wise group differences to assess the contribution of differences in the activation levels to distinct patterns between the two groups in each ROI.

#### Functional connectivity (FC) analyses

were next performed to investigate the contribution of the region(s) recruited specifically by the FHD+Typical children to the reading network during phonological processing using the CONN toolbox (Whitfield-Gabrieli & Nieto-Castanon, 2012). The preprocessed functional images were first band-pass filtered (0.008-0.09 Hz), detrended, and denoised to eliminate confounding effects of head movement and global hemodynamic changes using the anatomical aCompCor strategy (Behzadi et al., 2007; Chai et al., 2012). Task-relevant activation was also entered as a covariate of no interest to minimize the artificial interregional correlations caused by the experimental manipulations. Then, the timecourses specific to phonological processing were derived through weighting the residual time series by the task regressor specific to the FSM condition. The region(s) that showed activation preference for the FHD+Typical children, as identified from the whole-brain analyses, was applied as the seed region. Its timecourse was estimated using principal component analysis. Subject-wise FC maps were generated by correlating the timecourse of the seed region(s) with the timecourses of all remaining voxels, which were subsequently transformed to Fisher’s Z-scores. A left-hemispheric reading network, including the left inferior frontal cortex (pars opercularis and pars triangularis), left superior temporal gyrus, left inferior parietal cortex (inferior parietal lobule and supramarginal gyrus), and left fusiform gyrus was constructed following Preston et al. (2016) and applied as an explicit mask for second-level analyses. A two-sample comparison was performed to evaluate whether FHD+Typical and FHD-Typical children differed in the pre-reading functional connectivity strength between the seed region and left-hemispheric reading network. Significant regions were reported at a cluster-level of *p*_corrected_ < 0.05, Monte-Carlo corrected for multiple comparisons (voxel-level *p* < 0.005, *k* ≥ 50).

### DTI analyses

DTI data was analyzed in 31 typically developing children (14 FHD-Typical and 17 FHD+Typical). An established preprocessing protocol appropriate for this age range was applied (Wang et al., 2016). Specifically, a brain mask was first generated for each subject by removing the non-brain tissue from the corresponding T1 image using the Brain Extraction Tool (Smith, 2002) from Functional MRI of the Brain (FMRIB) software Library (Oxford, UK). Meanwhile, diffusion-weighted images collected in the DICOM format were converted into NRRD format using the DicomToNrrdConverter software (www.slicer.org). The DTIprep software was then applied to detect and correct for artifacts caused by eddy-currents, head motion, bed vibration and pulsation, venetian blind artifacts, as well as slice-wise and gradient-wise intensity inconsistencies (Oguz et al., 2014). Volumes with excessive motion defined as frame-wise head movement larger than 2 mm/0.5° were also identified and excluded from subsequent analyses. The two groups were not different in head movement for the remaining volumes (3 translational movement: left to right: *t*_29_ = 1.57, *p* = 0.13; posterior to anterior: *t*_29_ = 1.49, *p* = 0.15; bottom to top: *t*_29_ = 1.14, *p* = 0.26; 3 rotational movement: pitch: *t*_29_ = 1.94, *p* = 0.06; roll: *t*_29_ = 1.38, *p* = 0.18; yaw: *t*_29_ = 0.15, *p* = 0.88). The DTI images were further corrected for eddy currents and head motion using the VISTALab diffusion MRI software suite (www.vistalab.com). Diffusion tensors were then fitted using a linear least-squares fit, and FA maps were calculated for all subjects (Basser et al., 1994).

Fiber tractography was performed on the white matter tracts of interest using the Automated Fiber-tract Quantification (AFQ) toolbox (Yeatman et al., 2012). To do this, deterministic whole-brain streamline tractography was performed using an FA threshold of 0.2 and an angle threshold of 40°. Fibers were then segmented into separate tracts using two pre-defined anatomical ROIs (back projected from the MNI to the native space via T1 images) per tract as termination points. This was followed by a fiber-tract cleaning procedure to remove branch outliers from the core bundle. Each tract was then sampled to 100 equidistant nodes, and the diffusion property (i.e., the FA value in this case) for each node was estimated using a weighted mean of each fiber’s value based on its Mahalanobis distance from the fiber core. Finally, the 100 equidistant nodes of each tract were down-sampled to 50 nodes characterizing the central portion of the fiber tract. Following this method, the right superior longitudinal fasciculus (RSLF) was successfully identified in 31 children (14 FHD-Typical and 17 FHD+Typical), right inferior longitudinal fasciculus (RILF) in 30 participants (14 FHD-Typical and 16 FHD+Typical), and right arcuate fasciculus (RAF) in 18 subjects (9 FHD-Typical and 9 FHD+Typical). Due to the previously reported difficulties with reproducibility in defining the entire CC (Wakana et al., 2007), FA values were computed only for callosal fibers primarily linking bilateral occipital lobes (CC splenium) and (separately) those connecting frontal lobes (CC genu, see the supplementary video for a 3D render of the examined white matter tracts in one representative participant, codes adapted from https://github.com/yeatmanlab/AFQ/tree/master/3Dmesh). Tract reconstruction was successful in the CC genu for 28 children (12 FHD-Typical and 16 FHD+Typical) and in the CC splenium for 26 children (11 FHD-Typical and 15 FHD+Typical).

Two-sample t-tests were first carried out to evaluate FA differences between FHD+Typical and FHD-Typical children at each node in the four identified tracts. For subjects with both fMRI and DTI data available, Pearson-correlation analyses were further performed between functional activation level during phonological processing (i.e., the contrast estimate of FSM > VM) in the region(s) recruited by the FHD+Typical children and FA at each node in all the tracts (RSLF: 22 subjects, RILF: 22 subjects, RAF: 12 subjects, CC genu: 20 subjects, and CC splenium: 20 subjects). Significant results were reported at *p*_corrected_ < 0.05 for each node, FRD corrected for multiple comparisons.

## Results

### Longitudinal psychometric results (Table 1)

At the pre-reading stage, the FHD-Typical and FHD+Typical groups did not differ significantly from each other in terms of age (FHD-Typical: 65 ± 4.3 months; FHD+Typical: 66 ± 4.7 months, *t*_67_ = 1.34; *p* = 0.18). Both groups exhibited equivalent performance on measures of non-verbal IQ, early language competencies, letter knowledge, phonological processing, and rapid automatized naming abilities for objects and colors. Finally, no significant group differences were observed in terms of home literacy environment (HLE) and socio-economic status (SES), except that family members of FHD-Typical children reportedly read books, magazines or newspapers with the child more often than family members of FHD+Typical children (chi-square = 7.3, *p* = 0.03; see complete results on HLE and SES in Table S1 and Table S2, respectively).

All participants’ reading abilities were estimated at the emergent reading stage (between the end of 1^st^ and 4^th^ grade) after at least two years’ reading instruction. FHD+Typical and FHD-Typical children acquired equivalent scores in all four word-level reading assessments, including the WI and WA subtests of the WRMT-R, as well as the SWE and PDE subtests of the TOWRE. Moreover, to ensure that the latest time point captured a reliable estimation of reading performance along the developmental trajectory, comparative analyses was further conducted utilizing mean scores of each reading assessment across the performance of all the time points acquired. These analyses demonstrated comparable reading performance between the FHD-Typical and FHD+Typical children (see supplementary Table S3 for details), consistent with those based on the latest time point.

### Imaging results at the pre-reading stage

Consistent with the psychometric session, FHD-Typical and FHD+Typical children did not significantly differ in age during the scanning session (FHD-Typical: 66 ± 4.3 months; FHD+Typical: 68 ± 4.4 months, *t*_58_ = 1.8; *p* = 0.08).

### In-scanner task performance

Behavioral responses from four subjects (3 FHD-Typical and 1 FHD+Typical) could not be recorded due to technical issues. However, their imaging data was still included in analyses since high performance accuracies were demonstrated during the practice session and consistent button responses were observed by the accompanying research assistant during the formal experiment. Overall, no significant group differences were observed for subjects’ behavioral responses during this task (accuracy: *t*_54_ = 0.1, *p* = 0.92; reaction time: *t*_54_ = 1.4, *p* = 0.17, supplementary Table S4).

### FMRI results

#### Whole-brain results

Two-sample t-tests were first performed at the voxel level throughout the whole brain (i.e., mass-univariate analysis), but no significant differences were observed between the FHD-Typical and FHD+Typical groups (*p* < 0.05, FDR corrected). The whole-brain searchlight MVPA was subsequently performed, which identified two brain regions exhibiting distinct activation patterns between the FHD-Typical and FHD+Typical children in a combination of neighboring voxels (Figure 1A). The first region was in the right inferior frontal gyrus (RIFG) and comprised 5 connected searchlights which contained in 46 non-overlapping voxels spanning 1242 mm^3^ of volume (center-of-mass coordinate: [54,9,24]). The second one was in the left temporo-parietal cortex (LTPC), including 38 connected searchlights with 200 voxels occupying 5400 mm^3^ of volume (center-of-mass coordinate: [−45, −51, 24]).

**Figure 1.**
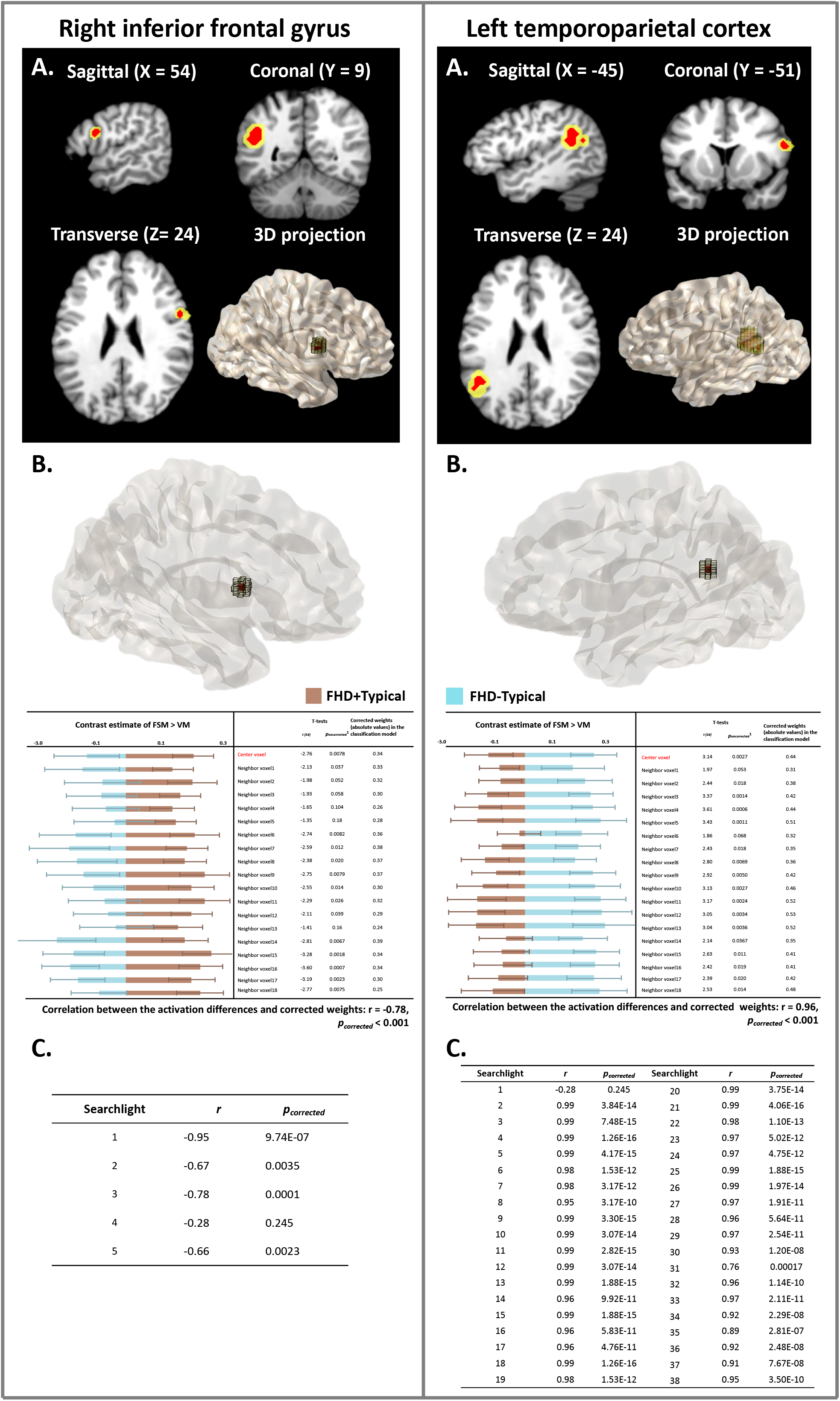
The right inferior frontal gyrus (RIFG, left section) and the left temporo-parietal cortex (LTPC, right section) exhibit distinct activation patterns between the FHD-Typical and FHD+Typical children, as revealed by the whole-brain searchlight MVPA. The top panel (A) shows RIFG and LTPC in slice-views and 3D projections; the significant regions are highlighted in yellow and the center voxels in red. The middle panel (B) illustrates that differences between the two groups of children in each voxel included in one example searchlight are significantly correlated with the contribution (corrected weight) of each of those voxels to the classification performance. Each representative searchlight is projected to a 3D image (center voxel in red). The bar figures below the images display the activation levels (contrast estimates of FSM > VM) for FHD+Typical (brown) and FHD-Typical children (blue) in each voxel. The tables show the statistical results of the group comparisons in each voxel and the absolute values of corrected weights in the classification model. The bottom panel (C) summarizes the correlation results for all significant searchlights. Whole-brain results are reported at *p*_corrected_ < 0.05, Bonferroni corrected for multiple comparisons. ^1.^ The results of the two-sample t-tests on the activation levels between the FHD+Typical and FHD-Typical groups in each voxel are not significant after FDR correction (*p*_corrected_ > 0.9)

Since a searchlight is a joint consideration of 19 neighboring voxels, follow-up analyses were conducted to understand how the group differences (FHD-Typical > FHD+Typical) in activation levels at each participating voxel contributed to the searchlight to trigger a significant group difference (Coutanche, 2013; Jimura & Poldrack, 2012). Across searchlights, larger group differences in the activation levels contributed more to the classification performance (i.e., higher absolute values of the corrected weights in the classification model, see example searchlights in the Figure 1B). Specifically, for the RIFG, all voxels showed higher activation levels for FHD+Typical than FHD-Typical children. Moreover, activation differences in the participating voxels were significantly and negatively correlated with the corrected weights (absolute values) derived from the classification models, in four of five searchlights (*r*_mean_ = −0.76, *p*_corrected_mean_ < 0.001, Figure 1C). This suggested that distinct activation patterns that reliably distinguished between the two groups in the RIFG region were mainly established on group differences in the FHD+Typical > FHD-Typical direction. By contrast, most of the voxels included in the LTPC searchlights showed higher activation levels for the FHD-Typical compared to the FHD+Typical children (98±8%), and the higher activation levels of FHD-Typical compared to FHD+Typical children were significantly correlated with the higher weights in the classification model in all but one of the thirty-eight searchlights (*r*_mean_ = 0.96, *p*_corrected_mean_ < 0.001, Figure 1C). This suggested that the significant group differences in the LTPC region were mainly based on group differences in the FHD-Typical > FHD+Typical direction, a pattern that was reverse to that in the RIFG region.

The same MVPA was further performed in each region as a whole, which rendered the same results. Distinct activation patterns were observed between the FHD-Typical and FHD+Typical children in both regions (RIFG: accuracy = 66.7%; *p_corrected_* < 0.006; LTPC: accuracy = 68.3%; *p_corrected_* < 0.02). Moreover, significant correlations were also observed between group differences in the activation levels and corrected weights extracted from the classification model for both regions (RIFG: *r* = −0.82; *p_corrected_* < 0.001; LTPC: r= 0.93; *p_corrected_* < 0.001).

#### Functional connectivity analyses

(FC, Figure 2) were further conducted using the RIFG as a seed, based on the higher activation levels identified in this region among FHD+Typical children. FC analyses revealed higher functional connectivity strength between the RIFG and the left inferior parietal cortex (LIPC), spanning over the left angular gyrus and the left inferior parietal lobule (([−36, −57, 42], k = 68 voxels) for the FHD+Typical compared to the FHD-Typical group. The opposite contrast did not reveal any significant functional pathway.

**Figure 2.**
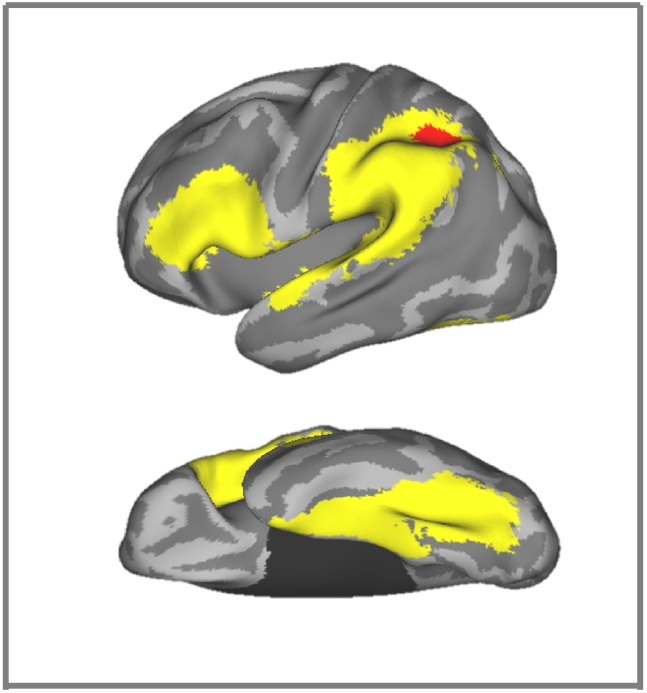
Sagittal (top) and transverse (bottom) views of the left hemisphere. Using the RIFG as the seed, functional connectivity (FC) analyses reveal stronger connectivity for FHD+Typical compared to FHD-Typical children in the left inferior parietal cortex (LIPC, highlighted in red) in a pre-defined reading mask (highlighted in yellow), including the inferior frontal cortex, temporo-parietal cortex (both in the sagittal view), and fusiform gyrus (transverse view). Results are reported at cluster-level *p*_corrected_ < 0.05, Monte-Carlo corrected for multiple comparisons (voxel-level *p* < 0.005, k ≥ 50).

### DTI and correlations with fMRI

Significant group differences were observed in bilateral segments of the CC splenium (nodes 2-4 and nodes 48-50, Figure 3), which demonstrated higher FA for FHD+Typical comparted to FHD-Typical children. No significant results were observed in other tracts (Supplementary Figure S1).

**Figure 3.**
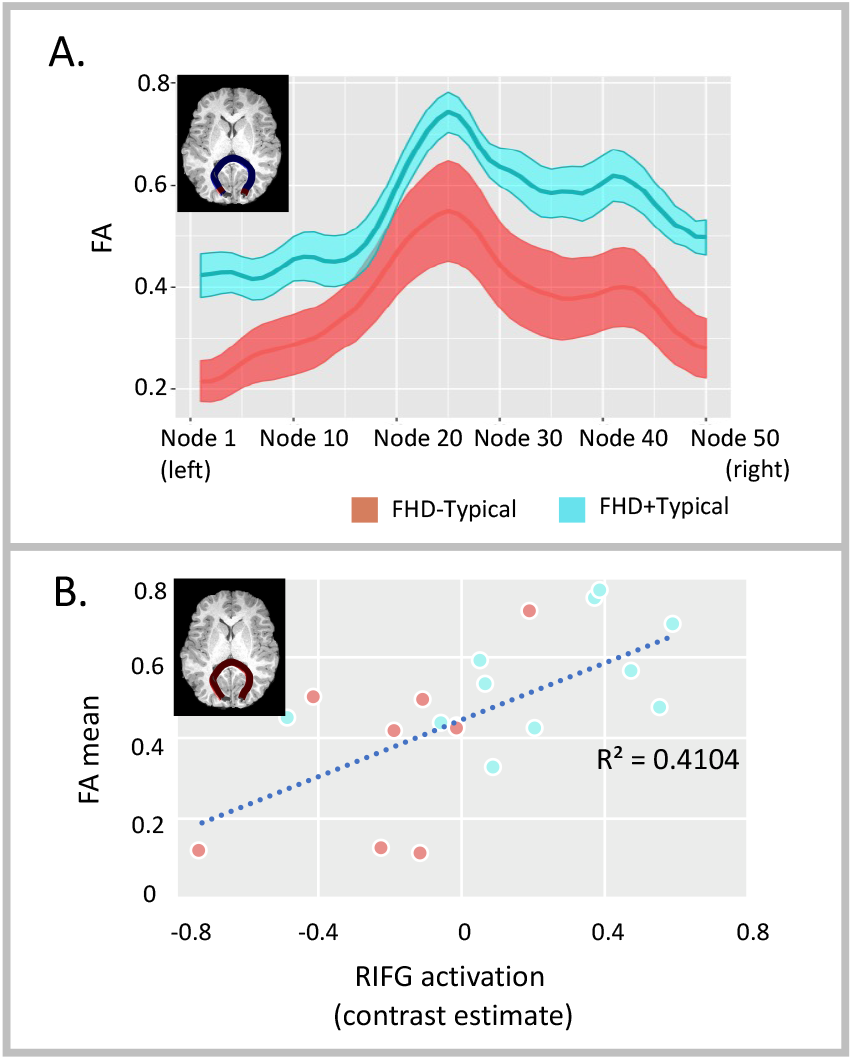
Panel A: Tract profiles (FA values at all 50 nodes) in the corpus callosum (CC) splenium for FHD-Typical (orange) and FHD+Typical (cyan) children. Two-sample t-tests reveal higher FA values in bilateral segments (nodes 2-4 and nodes 48-50, highlighted in red) in the CC splenium for the FHD+Typical compared to FHD-Typical children. Panel B: Correlation plot for mean FA in the corpus callosum splenium and activation level (contrast estimate of FSM > VM) in right inferior frontal gyrus during the phonological processing task. Significant positive correlations with activation levels in RIFG are observed at all nodes in the corpus callosum splenium.

Correlation analyses between RIFG activation level and FA values at all nodes in all the tracts demonstrated positive correlations at all nodes in the CC splenium (Figure 3C). The medial segment of the CC genu (nodes 22-28 and nodes 35-37) also exhibited significant correlations with RIFG activation level, but, these effects did not survive correction for multiple comparisons. No other significant correlations were observed.

## Discussion

The current study is the first to demonstrate the presence of putative neural protective mechanisms prior to reading onset in children who subsequently developed typical reading abilities despite a familial risk for dyslexia. Adopting a retrospective, longitudinal approach, FHD+ and FHD-children with typical reading abilities were characterized after at least two years of reading instruction. Through group comparisons of neural functional and structural (DTI) characteristics collected before the start of formal reading instruction, our results suggest an alternative network for phonological processing in FHD+Typical relative to FHD-Typical pre-readers. This was observed despite the fact that both groups subsequently developed equivalent typical reading abilities. Specifically, reduced activation was observed in the left temporo-parietal cortex (LTPC) in FHD+Typical relative to FHD-Typical children prior to reading onset (e.g., at the end of the pre-kindergarten/the beginning of kindergarten). This indicates that this neural characteristic is primarily associated with a familial risk rather than a diagnosis of developmental dyslexia. Meanwhile, a putative protective neural mechanism was revealed in FHD+Typical compared to FHD-Typical pre-readers, characterized by higher activations in the right inferior frontal gyrus (RIFG) and increased functional connectivity between RIFG and left inferior parietal cortex (LIPC). In addition, FHD+Typical compared to FHD-Typical children exhibited increased fractional anisotropy (FA) in the corpus callosum (CC), and the FA in the CC was positively correlated with RIFG activation. This suggests the development of an alternative anatomical infrastructure at the pre-reading stage which may then support a bilateral neural network for reading development in FHD+Typical children.

In the current analyses, the searchlight multivariate pattern analysis demonstrated that a combination of voxels can differentiate between FHD-Typical and FHD+Typical children despite no significant group differences observed in individual voxels. As illustrated in the example searchlights in Figure 1B, the neighboring voxels with weak differences in the activation levels (as demonstrated by *p*_corrected_ > 0.05) acted together in the searchlight and formed a strong classifier that significantly differentiated across groups. Further looking into the mechanism of how neighboring voxels acted together (Figure 1B and 1C) has revealed that significant classification results in the RIFG were mainly driven by voxels with higher activation in the FHD+Typical > FHD-Typical direction, while the performance in the LTPC were based on the voxel-wise activation in the FHD-Typical > FHD+Typical direction. Therefore, the MVPA exhibited subtle yet consistent and significant activation preferences for the FHD+Typical children in the RIFG and for the FHD-Typical children in the LTPC.

Hypoactivation in the LTPC was observed in FHD+Typical compared to FHD-Typical pre-readers despite their subsequent typical reading abilities, underscoring its role as a neural endophenotype associated with familial risk for dyslexia. Neural alterations in the LTPC have previously been reported in children with dyslexia (e.g., Peterson & Pennington, 2015; Richlan et al., 2009; Shaywitz & Shaywitz, 2008) as well as pre-reading children with a family history compared to typical controls (e.g., Im et al., 2015; Raschle et al., 2011, 2012, 2013). However, it remains unclear to what extent the observed neural differences may be attributed to the familial risk or represent as unique ‘premarker’ for developmental dyslexia. The current results demonstrate that the hypoactivation in the LTPC was present in FHD+ children who subsequently developed typical reading skills, and therefore strongly emphasize that this brain alteration is not a brain characteristic, or specific premarker, of dyslexia but rather seems to be an indication of atypical brain development associated with a familial risk. The observed link between LTPC deficits and FHD+ is further supported by recent genome-wide association studies which have shown significant correlations between variants in dyslexia susceptibility genes and both reading-related functional activation (Cope et al., 2012; Wilcke et al., 2012) and white matter volume (Darki et al., 2012) in LTPC. Due to the probabilistic nature of genetic transmission, it is possible that some FHD+Typical children might develop a typical, left-hemisphere-dominant reading network as a result of reduced/null genetic liability. However, for FHD+ children who show neural deficits in the LH reading network due to familial risk as observed in the current study, the development of compensatory/protective mechanisms seems to be important for acquiring typical reading skills.

Indeed, greater neural activation in RIFG and increased interhemispheric functional connectivity between the RIFG and LIPC was revealed in FHD+Typical compared to FHD-Typical children prior to reading onset, suggesting a potential protective mechanism. Increased activation in the RIFG has been previously associated with reading improvement in individuals with dyslexia and/or reading difficulties, therefore suggesting a compensatory mechanism, probably in response to successful intervention approaches (e.g., Eden et al., 2004; Farris et al., 2016; Hoeft et al., 2011; Richards et al., 2007; Temple et al., 2003). The current findings further add to the importance of the RIFG in typical reading development by demonstrating the reliance on RH frontal pathways in young FHD+ children (prereaders) who subsequently demonstrated successful reading development. The facilitative role of the RIFG was further corroborated by its functional connectivity to the left inferior parietal cortex, an area that has previously been shown to play an important role in reading development (e.g., Pugh et al., 2000; Schlaggar & McCandliss, 2007). Importantly, the RH pathway can already be observed in pre-readers before the start of any formal reading instruction/practice. This suggests that they may serve as a protective mechanism against adverse factors such as neural alternations in the LH reading network, and support the typical development of cognitive and pre-literacy (e.g., phonological processing) prerequisites for learning to read. This could reduce the likelihood of or even prevent children from developing reading impairments including developing dyslexia. It has been previously shown that structural connectivity precedes the development of the functional reading network (Saygin et al., 2016), suggesting that FHD+Typical children may show an alternative structural connectivity network very early in their language/literacy development (Langer et al., 2017) which then fosters the development of an alternative, protective reading network that enables typical reading development in FHD+Typical children.

Consistent with this conjecture, the RH protective pathways observed in FHD+Typical prereaders was further accompanied by enhanced cross-hemispheric structural connectivity in the Corpus Callosum (CC). Most of the CC neurons are excitatory (Fabri et al., 2014), and its importance for functional interhemispheric connectivity has been demonstrated numerous times (e.g., Cohen et al., 2000; Gooijers & Swinnen, 2014; Mohr et al., 1994). Greater FA in FHD+Typical compared to FHD-Typical children was only observed in the corpus callosum, but not in right-hemispheric white matter tracts which previously have been shown to be altered in compensated (elementary or middle school aged) readers (Hoeft et al., 2011; Wang et al., 2016). Higher FA values, specifically in the CC, in FHD+Typical children prior to reading onset might serve as a critical structural foundation that facilitates the recruitment of the right hemisphere during reading development. In support of this hypothesis, previous findings from our laboratory showed faster white matter development in RSLF for FHD+ children who subsequently develop into good readers compared to poor readers over the course of learning to read (Wang et al., 2016). Additionally, the observation of significant correlations between the microstructure of the CC and the neural activation in the RIFG during the phonological processing task are also in line with this notion. Interestingly, our interpretation is also in line with studies which examined the critical role the CC plays during literacy acquisition in adulthood, where an increased reliance on bi-hemispheric regions, most likely facilitated through the observed FA increases in the CC, was reported for adults who became literate in their twenties (Carreiras et al., 2009; Dehaene et al., 2015). Nevertheless, since information needs to be transferred via the CC to the right hemisphere, the observed protective/compensatory pathways might support typical reading development at the cost of speed. This speculation is consistent with the observation that FHD+Typical children in general read less fluently than FHD-Typical controls (e.g., Pennington & Lefly, 2001; Van Bergen et al., 2011). Future studies are needed to empirically evaluate the hypothesized association between reading fluency and a bilateral reading network.

Although not directly investigated in the current study, it is important to consider critical factors that might contribute to emergence of the protective/compensatory mechanisms in FHD+Typical children throughout early development. Investigation of brain characteristics in FHD+ children from an early age suggests a genetic role in shaping the hemispheric lateralization underlying language and reading development. Compared to a typical left-hemispheric dominance in FHD-controls, FHD+ prereaders and infants exhibit right-lateralized asymmetries in white matter tracts important for reading, as well as bilateral neural activation patterns in response to speech (Guttorm et al., 2001; Leppänen et al., 1999; Wang et al., 2016). The hypothesized genetic influences on atypical brain lateralization in FHD+ children are also in line with the Geschwind-Galaburda hypothesis of early development of atypical lateralization in individuals with dyslexia (Geschwind & Galaburda, 1985), and is further supported by the emerging findings suggesting that dyslexia-susceptibility genes are implicated in early brain development, such as cilia function, critical for subsequent hemispheric specialization (Brandler & Paracchini, 2014). Using genome-wide association techniques, dyslexia risk genes have also been directly associated with atypical development of the CC and hemispheric lateralization (Darki et al., 2012; Pinel et al., 2012). Based on these findings, it can be hypothesized that children with a familial risk of dyslexia are genetically predisposed for a bilateral neural mechanism underlying reading development, setting a foundation for the development of RH protective/compensatory functional networks.

Moreover, several environmental factors have also been identified to facilitate the formation of the protective/compensatory functional pathways in the right hemisphere. For example, enriched early home literacy and higher SES have been shown to associated with enhanced recruitment of right-hemispheric perisylvian and frontal areas for language and reading processing in children (Noble et al., 2006; Powers et al., 2016). Importantly, the association between HLE and right-hemispheric activation was specific for FHD+ but not FHD-pre-readers (Powers et al., 2016), suggesting a specific gene x environment interaction in the right hemisphere in FHD+ children. In addition to family characteristics, although debated, educational experiences such as music training and specialized teaching strategies have also been shown to shape the neural mechanisms underlying reading development towards a bilateral network accompanied with stronger interhemispheric structural microstructure of CC (Habibi et al., 2016; Mei et al., 2013; Yoncheva et al., 2010; 2015; Zuk et al., 2018). Altogether, one can postulate that positive environmental stimulation, such as enriched home literacy environment, interacts with genetic predisposition and collectively they enable the development of protective/compensatory neural mechanisms in the right hemisphere to mediate the deficient processing in the LH and through this support typical reading development in FHD+Typical children (Yu et al., 2018).

The presence of distinct neural characteristics in FHD+Typical pre-readers compared to controls also provides valuable insight into optimal early diagnosis and intervention approaches. Both neural deficits and putative protective mechanisms were observed in the FHD+Typical children at the prereading stage, encouraging a holistic approach that considers both risk and protective aspects when screening for early risk of reading impairment as well as for designing early intervention programs for children at risk for dyslexia (Ozernov-Palchik et al., 2016). Moreover, the establishment of protective pathways prior to reading onset also opens the possibility of developing resilience in at-risk children through preventative intervention approaches administered at earlier stages during the course of reading development. This may result in children experiencing reduced learning difficulties while learning to read or even exhibiting typical reading development, as observed in the FHD+Typical children.

This study provides the first evidence for the development of putative protective neural mechanisms in FHD+ pre-readers who subsequently develop typical reading skills, but it is to be interpreted in the context of two considerations. First, the data from FHD+ children who subsequently showed reading impairments (i.e., FHD+Impaired) were not included in our current analysis due to the small sample size (n=12) available in this longitudinal data set. Therefore, it is unknown whether any neural protective pathways were available to support reading development, albeit not successful, in FHD+Impaired children. Nevertheless, it is interesting to note that the putative protective neural mechanisms observed in the current study for the FHD+Typical pre-readers are similar to those that emerged in school-age children with dyslexia after successful intervention (e.g., Horowitz-Kraus et al., 2014; Temple et al., 2003), suggesting a protracted time window for the development of protective/compensatory neural mechanisms possibly driven by multiple factors. Future studies with a large sample are needed to determine whether, and if so when/how, compensatory/protective mechanisms interact with risk factors and collectively contribute to reading outcomes among FHD+ children. Second, the genetic contributions for the development of protective pathways were only tested indirectly in this study, since only children with a reported family history of dyslexia were examined. Future longitudinal studies ranging from infancy to school age are needed and both genetic and environmental measures should be considered. These studies will help to identify how early in a child’s life these protective neural mechanisms emerge (e.g. are they present at birth or develop over time) and what genetic and environmental (e.g. home literacy, quality of language input) factors facilitate their emergence over the developmental time course. Answering these research questions could inform the design of preventative and remediation strategies for children at risk for dyslexia.

## Conclusion

Despite an increased risk of developing dyslexia, about half of children with a familial risk for dyslexia develop typical reading skills. The current study is the first to demonstrate that putative neural protective mechanisms that seem to support typical reading abilities in children with a familial risk who subsequently develop typical reading skills are present before the onset of formal reading instruction. Specifically, FHD+Typical pre-readers exhibited higher activation in the RIFG during phonological processing compared to FHD-Typical children, which was further supported by increased interhemispheric functional and structural connectivity. The current findings support a working hypothesis of potential neural protective and compensatory pathways in FHD+Typical children which emerge through interactions between genetics, neurobiology, and environmental factors to facilitate their typical reading development. Future longitudinal studies are needed to explore the factors, both genetic and environmental, that enable these putative protective and compensatory mechanisms, as well as their developmental trajectories. Such research may guide the design of early assessment and interventional tools for children at risk for dyslexia.

## References

Aboitiz, F., & Montiel, J. (2003). One hundred million years of interhemispheric communication: the history of the corpus callosum. Brazilian journal of medical and biological research, 36(4), 409–420.

Ashburner, J. (2007). A fast diffeomorphic image registration algorithm. Neuroimage, 38(1), 95–113.

Astrom, R. L., Wadsworth, S. J., & DeFries, J. C. (2007). Etiology of the stability of reading difficulties: the longitudinal twin study of reading disabilities. Twin Research and Human Genetics, 10(3), 434–439. doi:https://doi.org/10.1375/twin.10.3.434

Baker, S. F., & Ireland, J. L. (2007). The link between dyslexic traits, executive functioning, impulsivity and social self-esteem among an offender and non-offender sample. International Journal of Law and Psychiatry, 30(6), 492–503.

Barquero, L. A., Davis, N., & Cutting, L. E. (2014). Neuroimaging of reading intervention: a systematic review and activation likelihood estimate meta-analysis. PloS one, 9(1), e83668. doi:https://doi.org/10.1371/journal.pone.0083668

Basser, P. J., Mattiello, J., & LeBihan, D. (1994). MR diffusion tensor spectroscopy and imaging. Biophysical journal, 66(1), 259–267.

Bauer, A. J., & Just, M. A. (2017). A brain-based account of “basic-level” concepts. Neuroimage, 161, 196–205.

Behzadi, Y., Restom, K., Liau, J., & Liu, T. T. (2007). A component based noise correction method (CompCor) for BOLD and perfusion based fMRI. Neuroimage, 37(1), 90–101.

Boets, B., Wouters, J., Van Wieringen, A., & Ghesquiere, P. (2007). Auditory processing, speech perception and phonological ability in pre-school children at high-risk for dyslexia: a longitudinal study of the auditory temporal processing theory. Neuropsychologia, 45(8), 1608–1620.

Brandler, W. M., & Paracchini, S. (2014). The genetic relationship between handedness and neurodevelopmental disorders. Trends in Molecular Medicine, 20(2), 83–90.

Carreiras, M., Seghier, M. L., Baquero, S., Estévez, A., Lozano, A., Devlin, J. T., & Price, C. J. (2009). An anatomical signature for literacy. Nature, 461(7266), 983.

Chai, X. J., Castañón, A. N., Öngür, D., & Whitfield-Gabrieli, S. (2012). Anticorrelations in resting state networks without global signal regression. Neuroimage, 59(2), 1420–1428.

Cohen, L., Dehaene, S., Naccache, L., Lehéricy, S., Dehaene-Lambertz, G., Hénaff, M.-A., & Michel, F. (2000). The visual word form area: spatial and temporal characterization of an initial stage of reading in normal subjects and posterior split-brain patients. Brain, 123(2), 291–307.

Cope, N., Eicher, J. D., Meng, H., Gibson, C. J., Hager, K., Lacadie, C.,… Gruen, J. R. (2012). Variants in the DYX2 locus are associated with altered brain activation in reading-related brain regions in subjects with reading disability. Neuroimage, 63(1), 148–156.

Coutanche, M. N. (2013). Distinguishing multi-voxel patterns and mean activation: why, how, and what does it tell us? Cognitive, Affective, & Behavioral Neuroscience, 13(3), 667–673.

Darki, F., Peyrard-Janvid, M., Matsson, H., Kere, J., & Klingberg, T. (2012). Three dyslexia susceptibility genes, DYX1C1, DCDC2, and KIAA0319, affect temporo-parietal white matter structure. Biological psychiatry, 72(8), 671–676.

Dehaene, S., Cohen, L., Morais, J., & Kolinsky, R. (2015). Illiterate to literate: behavioural and cerebral changes induced by reading acquisition. Nature Reviews Neuroscience, 16(4), 234.

Denney, M. K., English, J. P., Gerber, M. M., Leafstedt, J., & Ruz, M. L. (2001). Family and Home Literacy Practices: Mediating Factors for Preliterate English Learners at Risk.

Dougherty, C. (2003). Numeracy, literacy and earnings: evidence from the National Longitudinal Survey of Youth. Economics of education review, 22(5), 511–521.

Eden, G. F., Jones, K. M., Cappell, K., Gareau, L., Wood, F. B., Zeffiro, T. A.,… Flowers, D. L. (2004). Neural changes following remediation in adult developmental dyslexia. Neuron, 44(3), 411–422.

Evans, A. C. (1992). An MRI-based stereotactic atlas from 250 young normal subjects. Soc. neurosci. abstr, 1992.

Fabri, M., Pierpaoli, C., Barbaresi, P., & Polonara, G. (2014). Functional topography of the corpus callosum investigated by DTI and fMRI. World journal of radiology, 6(12), 895.

Farris, E. A., Ring, J., Black, J., Lyon, G. R., & Odegard, T. N. (2016). Predicting Growth in Word Level Reading Skills in Children With Developmental Dyslexia Using an Object Rhyming Functional Neuroimaging Task. Developmental neuropsychology, 41(3), 145–161.

Friston, K. J., Holmes, A. P., Worsley, K. J., Poline, J. P., Frith, C. D., & Frackowiak, R. S. (1994). Statistical parametric maps in functional imaging: a general linear approach. Human Brain Mapping, 2(4), 189–210.

Gaab, N., Gabrieli, J. D., & Glover, G. H. (2007a). Assessing the influence of scanner background noise on auditory processing. I. An fMRI study comparing three experimental designs with varying degrees of scanner noise. Human Brain Mapping, 28(8), 703–720.

Gaab, N., Gabrieli, J. D., & Glover, G. H. (2007b). Assessing the influence of scanner background noise on auditory processing. II. An fMRI study comparing auditory processing in the absence and presence of recorded scanner noise using a sparse design. Human Brain Mapping, 28(8), 721–732.

Gallagher, A., Frith, U., & Snowling, M. J. (2000). Precursors of literacy delay among children at genetic risk of dyslexia. The Journal of Child Psychology and Psychiatry and Allied Disciplines, 41(2), 203–213.

Geschwind, N., & Galaburda, A. M. (1985). Cerebral lateralization: Biological mechanisms, associations, and pathology: I. A hypothesis and a program for research. Archives of neurology, 42(5), 428–459.

Gooijers, J., & Swinnen, S. (2014). Interactions between brain structure and behavior: the corpus callosum and bimanual coordination. Neuroscience & Biobehavioral Reviews, 43, 1–19.

Guttorm, T. K., Leppänen, P. H., Richardson, U., & Lyytinen, H. (2001). Event-related potentials and consonant differentiation in newborns with familial risk for dyslexia. Journal of Learning Disabilities, 34(6), 534–544.

Habibi, A., Cahn, B. R., Damasio, A., & Damasio, H. (2016). Neural correlates of accelerated auditory processing in children engaged in music training. Developmental cognitive neuroscience, 21, 1–14.

Haft, S. L., Myers, C. A., & Hoeft, F. (2016). Socio-emotional and cognitive resilience in children with reading disabilities. Current opinion in behavioral sciences, 10, 133–141.

Haufe, S., Meinecke, F., Görgen, K., Dähne, S., Haynes, J.-D., Blankertz, B., & Bießmann, F. (2014). On the interpretation of weight vectors of linear models in multivariate neuroimaging. Neuroimage, 87, 96–110.

Haxby, J. V., Gobbini, M. I., Furey, M. L., Ishai, A., Schouten, J. L., & Pietrini, P. (2001). Distributed and overlapping representations of faces and objects in ventral temporal cortex. Science, 293(5539), 2425–2430.

Hebart, M. N., Görgen, K., & Haynes, J.-D. (2015). The Decoding Toolbox (TDT): a versatile software package for multivariate analyses of functional imaging data. Frontiers in neuroinformatics, 8, 88.

Hinkley, L. B., Marco, E. J., Brown, E. G., Bukshpun, P., Gold, J., Hill, S.,… Barkovich, A. J. (2016). The contribution of the corpus callosum to language lateralization. Journal of Neuroscience, 36(16), 4522–4533.

Hoeft, F., McCandliss, B. D., Black, J. M., Gantman, A., Zakerani, N., Hulme, C.,… Reiss, A. L. (2011). Neural systems predicting long-term outcome in dyslexia. Proceedings of the National Academy of Sciences, 108(1), 361–366.

Horowitz-Kraus, T., Wang, Y., Plante, E., & Holland, S. K. (2014). Involvement of the right hemisphere in reading comprehension: a DTI study. Brain Research, 1582, 34–44.

Im, K., Raschle, N. M., Smith, S. A., Ellen Grant, P., & Gaab, N. (2015). Atypical sulcal pattern in children with developmental dyslexia and at-risk kindergarteners. Cerebral Cortex, 26(3), 1138–1148.

International Dyslexia Association (IDA, 2007). Dyslexia basics. Retrieved October, 21, 2007.

Jimura, K., & Poldrack, R. A. (2012). Analyses of regional-average activation and multivoxel pattern information tell complementary stories. Neuropsychologia, 50(4), 544–552.

Katusic, S. K., Colligan, R. C., Barbaresi, W. J., Schaid, D. J., & Jacobsen, S. J. (2001). Incidence of reading disability in a population-based birth cohort, 1976–1982, Rochester, Minn. Paper presented at the Mayo Clinic Proceedings.

Kaufman, A., & Kaufman, N. (2004). Brief Intelligence Test. 2. [Press release]

Koster, C., Been, P. H., Krikhaar, E. M., Zwarts, F., Diepstra, H. D., & Van Leeuwen, T. H. (2005). Differences at 17 months: Productive language patterns in infants at familial risk for dyslexia and typically developing infants. Journal of Speech, Language, and Hearing Research, 48(2), 426–438.

Kragel, P. A., Carter, R. M., & Huettel, S. A. (2012). What makes a pattern? Matching decoding methods to data in multivariate pattern analysis. Frontiers in neuroscience, 6, 162.

Kriegeskorte, N., Goebel, R., & Bandettini, P. (2006). Information-based functional brain mapping. Proceedings of the National Academy of Sciences, 103(10), 3863–3868.

Langer, N., Peysakhovich, B., Zuk, J., Drottar, M., Sliva, D. D., Smith, S.,… Gaab, N. (2017). White matter alterations in infants at risk for developmental dyslexia. Cerebral Cortex, 27(2), 1027–1036.

Leppänen, P. H., Pihko, E., Eklund, K. M., & Lyytinen, H. (1999). Cortical responses of infants with and without a genetic risk for dyslexia: II. Group effects. Neuroreport, 10(5), 969–973.

Lohvansuu, K., Hämäläinen, J. A., Tanskanen, A., Ervast, L., Heikkinen, E., Lyytinen, H., & Leppänen, P. H. (2014). Enhancement of brain event-related potentials to speech sounds is associated with compensated reading skills in dyslexic children with familial risk for dyslexia. International journal of psychophysiology, 94(3), 298–310.

Lyytinen, H., Ahonen, T., Eklund, K., Guttorm, T., Kulju, P., Laakso, M. L.,… Poikkeus, A. M. (2004). Early development of children at familial risk for Dyslexia—follow-up from birth to school age. Dyslexia, 10(3), 146–178.

Lyytinen, H., Ahonen, T., Eklund, K., Guttorm, T. K., Laakso, M.-L., Leinonen, S.,… Puolakanaho, A. (2001). Developmental pathways of children with and without familial risk for dyslexia during the first years of life. Developmental neuropsychology, 20(2), 535–554.

Lyytinen, H., Guttorm, T. K., Huttunen, T., Hämäläinen, J., Leppänen, P. H., & Vesterinen, M. (2005). Psychophysiology of developmental dyslexia: A review of findings including studies of children at risk for dyslexia. Journal of Neurolinguistics, 18(2), 167–195.

Maurer, U., Bucher, K., Brem, S., & Brandeis, D. (2003). Altered responses to tone and phoneme mismatch in kindergartners at familial dyslexia risk. Neuroreport, 14(17), 2245–2250.

McCandliss, B. D., & Noble, K. G. (2003). The development of reading impairment: a cognitive neuroscience model. Mental retardation and developmental disabilities research reviews, 9(3), 196–205.

Mei, L., Xue, G., Lu, Z.-L., He, Q., Zhang, M., Xue, F.,… Dong, Q. (2013). Orthographic transparency modulates the functional asymmetry in the fusiform cortex: An artificial language training study. Brain and language, 125(2), 165–172.

Mohr, B., Pulvermüller, F., Rayman, J., & Zaidel, E. (1994). Interhemispheric cooperation during lexical processing is mediated by the corpus callosum: Evidence from the split-brain. Neuroscience letters, 181(1-2), 17–21.

Morgan, P. L., Fuchs, D., Compton, D. L., Cordray, D. S., & Fuchs, L. S. (2008). Does early reading failure decrease children’s reading motivation? Journal of Learning Disabilities, 41(5), 387–404.

Newbury, D., Paracchini, S., Scerri, T., Winchester, L., Addis, L., Richardson, A. Monaco, A. (2011). Investigation of dyslexia and SLI risk variants in reading-and language-impaired subjects. Behavior genetics, 41(1), 90–104.

Noble, K. G., Wolmetz, M. E., Ochs, L. G., Farah, M. J., & McCandliss, B. D. (2006). Brain–behavior relationships in reading acquisition are modulated by socioeconomic factors. Developmental science, 9(6), 642–654.

Norman, K. A., Polyn, S. M., Detre, G. J., & Haxby, J. V. (2006). Beyond mind-reading: multi-voxel pattern analysis of fMRI data. Trends in cognitive sciences, 10(9), 424–430.

Norton, E. S., Beach, S. D., & Gabrieli, J. D. (2015). Neurobiology of dyslexia. Current opinion in neurobiology, 30, 73–78.

Oguz, I., Farzinfar, M., Matsui, J., Budin, F., Liu, Z., Gerig, G.,… Styner, M. A. (2014). DTIPrep: quality control of diffusion-weighted images. Frontiers in neuroinformatics, 8, 4.

Ozernov-Palchik, O., Yu, X., Wang, Y., & Gaab, N. (2016). Lessons to be learned: how a comprehensive neurobiological framework of atypical reading development can inform educational practice. Current opinion in behavioral sciences, 10, 45–58.

Ozernov-Palchik, O., & Gaab, N. (2016). Tackling the ‘dyslexia paradox’: reading brain and behavior for early markers of developmental dyslexia. Wiley Interdisciplinary Reviews: Cognitive Science, 7(2), 156–176. doi:http://dx.doi.org/10.1002/wcs.1383

Pennington, B. F., & Lefly, D. L. (2001). Early reading development in children at family risk for dyslexia. Child development, 72(3), 816–833.

Peterson, R. L., & Pennington, B. F. (2015). Developmental dyslexia. Annual review of clinical psychology, 11, 283–307.

Pinel, P., Fauchereau, F., Moreno, A., Barbot, A., Lathrop, M., Zelenika, D.,… Dehaene, S. (2012). Genetic variants of FOXP2 and KIAA0319/TTRAP/THEM2 locus are associated with altered brain activation in distinct language-related regions. Journal of Neuroscience, 32(3), 817–825.

Plakas, A., van Zuijen, T., van Leeuwen, T., Thomson, J. M., & van der Leij, A. (2013). Impaired non-speech auditory processing at a pre-reading age is a risk-factor for dyslexia but not a predictor: an ERP study. Cortex, 49(4), 1034–1045.

Poelmans, G., Buitelaar, J., Pauls, D., & Franke, B. (2011). A theoretical molecular network for dyslexia: integrating available genetic findings. Molecular psychiatry, 16(4), 365–382.

Powers, S. J., Wang, Y., Beach, S. D., Sideridis, G. D., & Gaab, N. (2016). Examining the relationship between home literacy environment and neural correlates of phonological processing in beginning readers with and without a familial risk for dyslexia: an fMRI study. Annals of dyslexia, 66(3), 337–360.

Pugh, K. R., Mencl, W. E., Jenner, A. R., Katz, L., Frost, S. J., Lee, J. R.,… Shaywitz, B. A. (2000). Functional neuroimaging studies of reading and reading disability (developmental dyslexia). Mental retardation and developmental disabilities research reviews, 6(3), 207–213.

Rajapakse, J. C., Giedd, J. N., & Rapoport, J. L. (1997). Statistical approach to segmentation of single-channel cerebral MR images. IEEE transactions on medical imaging, 16(2), 176–186.

Raschle, N., Zuk, J., Ortiz-Mantilla, S., Sliva, D. D., Franceschi, A., Grant, P. E.,… Gaab, N. (2012). Pediatric neuroimaging in early childhood and infancy: challenges and practical guidelines. Annals of the New York Academy of sciences, 1252(1), 43–50.

Raschle, N. M., Chang, M., & Gaab, N. (2011). Structural brain alterations associated with dyslexia predate reading onset. Neuroimage, 57(3), 742–749.

Raschle, N. M., Lee, M., Buechler, R., Christodoulou, J. A., Chang, M., Vakil, M.,… Gaab, N. (2009). Making MR imaging child’s play-pediatric neuroimaging protocol, guidelines and procedure. Journal of visualized experiments: JoVE(29).

Raschle, N. M., Stering, P. L., Meissner, S. N., & Gaab, N. (2013). Altered neuronal response during rapid auditory processing and its relation to phonological processing in prereading children at familial risk for dyslexia. Cerebral Cortex, 24(9), 2489–2501.

Raschle, N. M., Zuk, J., & Gaab, N. (2012). Functional characteristics of developmental dyslexia in left-hemispheric posterior brain regions predate reading onset. Proceedings of the National Academy of Sciences, 109(6), 2156–2161.

Richards, T., Berninger, V., Winn, W., Stock, P., Wagner, R., Muse, A., & Maravilla, K. (2007). Functional MRI activation in children with and without dyslexia during pseudoword aural repeat and visual decode: Before and after treatment. Neuropsychology, 21(6), 732.

Richlan, F., Kronbichler, M., & Wimmer, H. (2009). Functional abnormalities in the dyslexic brain: A quantitative meta-analysis of neuroimaging studies. Human Brain Mapping, 30(10), 3299–3308.

Richlan, F., Kronbichler, M., & Wimmer, H. (2011). Meta-analyzing brain dysfunctions in dyslexic children and adults. Neuroimage, 56(3), 1735–1742. doi:https://doi.org/10.1016/j.neuroimage.2011.02.040

Richlan, F., Kronbichler, M., & Wimmer, H. (2013). Structural abnormalities in the dyslexic brain: a metaanalysis of voxel-based morphometry studies. Human Brain Mapping, 34(11), 3055–3065. doi:http://dx.doi.org/10.1002/hbm.22127

Saygin, Z. M., Osher, D. E., Norton, E. S., Youssoufian, D. A., Beach, S. D., Feather, J.,… Kanwisher, N. (2016). Connectivity precedes function in the development of the visual word form area. Nature neuroscience, 19(9), 1250.

Scarborough, H. S. (1990). Very early language deficits in dyslexic children. Child development, 61(6), 1728–1743.

Scerri, T. S., Darki, F., Newbury, D. F., Whitehouse, A. J., Peyrard-Janvid, M., Matsson, H.,… Stein, J. (2012). The dyslexia candidate locus on 2p12 is associated with general cognitive ability and white matter structure. PloS one, 7(11), e50321.

Scerri, T. S., Morris, A. P., Buckingham, L.-L., Newbury, D. F., Miller, L. L., Monaco, A. P.,… Paracchini, S. (2011). DCDC2, KIAA0319 and CMIP are associated with reading-related traits. Biological psychiatry, 70(3), 237–245.

Schlaggar, B. L., & McCandliss, B. D. (2007). Development of neural systems for reading. Annu. Rev. Neurosci., 30, 475–503. doi:https://doi.org/10.1146/annurev.neuro.28.061604.135645

Semel, E., Wiig, E., & Secord, W. (2003). CELF 4: Clinical Evaluation of Language Fundamentals (4th Ed.) [Press release]

Shaywitz, S. E., & Shaywitz, B. A. (2008). Paying attention to reading: the neurobiology of reading and dyslexia. Development and psychopathology, 20(4), 1329–1349.

Shaywitz, S. E., Shaywitz, B. A., Fulbright, R. K., Skudlarski, P., Mencl, W. E., Constable, R. T.,… Fletcher, J. M. (2003). Neural systems for compensation and persistence: young adult outcome of childhood reading disability. Biological psychiatry, 54(1), 25–33.

Smith, S. M. (2002). Fast robust automated brain extraction. Human Brain Mapping, 17(3), 143–155.

Snowling, M. J., Gallagher, A., & Frith, U. (2003). Family risk of dyslexia is continuous: Individual differences in the precursors of reading skill. Child development, 74(2), 358–373.

Snowling, M. J., & Melby-Lervåg, M. (2016). Oral language deficits in familial dyslexia: A meta-analysis and review. Psychological bulletin, 142(5), 498. doi:http://dx.doi.org/10.1037/bul0000037

Stelzer, J., Chen, Y., & Turner, R. (2013). Statistical inference and multiple testing correction in classification-based multi-voxel pattern analysis (MVPA): random permutations and cluster size control. Neuroimage, 65, 69–82.

Taipale, M., Kaminen, N., Nopola-Hemmi, J., Haltia, T., Myllyluoma, B., Lyytinen, H.,… Hannula-Jouppi, K. (2003). A candidate gene for developmental dyslexia encodes a nuclear tetratricopeptide repeat domain protein dynamically regulated in brain. Proceedings of the National Academy of Sciences, 100(20), 11553–11558.

Temple, E., Deutsch, G. K., Poldrack, R. A., Miller, S. L., Tallal, P., Merzenich, M. M., & Gabrieli, J. D. (2003). Neural deficits in children with dyslexia ameliorated by behavioral remediation: evidence from functional MRI. Proceedings of the National Academy of Sciences, 100(5), 2860–2865.

Torgesen, J. K., Rashotte, C. A., & Wagner, R. K. (1999). TOWRE: Test of word reading efficiency: Pro-ed Austin, TX.

Torppa, M., Lyytinen, P., Erskine, J., Eklund, K., & Lyytinen, H. (2010). Language development, literacy skills, and predictive connections to reading in Finnish children with and without familial risk for dyslexia. Journal of Learning Disabilities, 43(4), 308–321. doi:https://doi.org/10.1177/0022219410369096

Tzourio-Mazoyer, N., Landeau, B., Papathanassiou, D., Crivello, F., Etard, O., Delcroix, N.,… Joliot, M. (2002). Automated anatomical labeling of activations in SPM using a macroscopic anatomical parcellation of the MNI MRI single-subject brain. Neuroimage, 15(1), 273–289.

Van Bergen, E., De Jong, P. F., Plakas, A., Maassen, B., & van der Leij, A. (2012). Child and parental literacy levels within families with a history of dyslexia. Journal of Child Psychology and Psychiatry, 53(1), 28–36.

Van Bergen, E., De Jong, P. F., Regtvoort, A., Oort, F., Van Otterloo, S., & van der Leij, A. (2011). Dutch children at family risk of dyslexia: Precursors, reading development, and parental effects. Dyslexia, 17(1), 2–18.

van der Leij, A., Van Bergen, E., van Zuijen, T., De Jong, P., Maurits, N., & Maassen, B. (2013). Precursors of developmental dyslexia: an overview of the longitudinal Dutch dyslexia programme study. Dyslexia, 19(4), 191–213.

van Herten, M., Pasman, J., van Leeuwen, T. H., Been, P. H., van der Leij, A., Zwarts, F., & Maassen, B. (2008). Differences in AERP responses and atypical hemispheric specialization in 17-month-old children at risk of dyslexia. Brain Research, 1201, 100–105.

van Leeuwen, T., Been, P., Kuijpers, C., Zwarts, F., Maassen, B., & van der Leij, A. (2006). Mismatch response is absent in 2-month-old infants at risk for dyslexia. Neuroreport, 17(4), 351–355.

van Leeuwen, T., Been, P., van Herten, M., Zwarts, F., Maassen, B., & van der Leij, A. (2007). Cortical categorization failure in 2-month-old infants at risk for dyslexia. Neuroreport, 18(9), 857–861.

Vandermosten, M., Vanderauwera, J., Theys, C., De Vos, A., Vanvooren, S., Sunaert, S.,… Ghesquière, P. (2015). A DTI tractography study in pre-readers at risk for dyslexia. Developmental cognitive neuroscience, 14, 8–15.

Wagner, R., Torgesen, J., Rashotte, C., & Pearson, N. A. (1999). CTOPP: Comprehensive Test of Phonological Processing–Second Edition.

Wakana, S., Caprihan, A., Panzenboeck, M. M., Fallon, J. H., Perry, M., Gollub, R. L.,… Dubey, P. (2007). Reproducibility of quantitative tractography methods applied to cerebral white matter. Neuroimage, 36(3), 630–644.

Wang, Y., Mauer, M. V., Raney, T., Peysakhovich, B., Becker, B. L., Sliva, D. D., & Gaab, N. (2016). Development of tract-specific white matter pathways during early reading development in at-risk children and typical controls. Cerebral Cortex, 27(4), 2469–2485.

Whitfield-Gabrieli, S., & Nieto-Castanon, A. (2012). Conn: a functional connectivity toolbox for correlated and anticorrelated brain networks. Brain connectivity, 2(3), 125–141.

Wilcke, A., Ligges, C., Burkhardt, J., Alexander, M., Wolf, C., Quente, E.,… Müller-Myhsok, B. (2012). Imaging genetics of FOXP2 in dyslexia. European Journal of Human Genetics, 20(2), 224.

Wilke, M., Holland, S. K., Altaye, M., & Gaser, C. (2008). Template-O-Matic: a toolbox for creating customized pediatric templates. Neuroimage, 41(3), 903–913.

Wolf, M., & Denckla, M. B. (2005). RAN/RAS: Rapid automatized naming and rapid alternating stimulus tests: Pro-ed Austin, TX.

Woodcock, R. W. (1987). Woodcock reading mastery tests, revised: American Guidance Service Circle Pines, MN.

Yeatman, J. D., Dougherty, R. F., Myall, N. J., Wandell, B. A., & Feldman, H. M. (2012). Tract profiles of white matter properties: automating fiber-tract quantification. PloS one, 7(11), e49790.

Yoncheva, Y. N., Blau, V. C., Maurer, U., & McCandliss, B. D. (2010). Attentional focus during learning impacts N170 ERP responses to an artificial script. Developmental neuropsychology, 35(4), 423–445.

Yoncheva, Y. N., Wise, J., & McCandliss, B. (2015). Hemispheric specialization for visual words is shaped by attention to sublexical units during initial learning. Brain and language, 145, 23–33. doi:https://doi.org/10.1016/j.bandl.2015.04.001

Yu, X., Raney, T., Perdue, M. V., Zuk, J., Ozernov-Palchik, O., Becker, B. L.,… Gaab, N. (2018). Emergence of the neural network underlying phonological processing from the prereading to the emergent reading stage: A longitudinal study. Human Brain Mapping, 39(5), 2047–2063.

Yu, X., Zuk, J., & Gaab, N. (2018). What Factors Facilitate Resilience in Developmental Dyslexia? Examining Protective and Compensatory Mechanisms Across the Neurodevelopmental Trajectory. Child development perspectives.

Zuk, J., Perdue, M. V., Becker, B., Yu, X., Chang, M., Raschle, N. M., & Gaab, N. (2018). Neural correlates of phonological processing: Disrupted in children with dyslexia and enhanced in musically trained children. Developmental cognitive neuroscience, 34, 82–91.

