## Supplementary materials for "Putative protective neural mechanisms in pre-readers with a family history of dyslexia who subsequently develop typical reading skills"

**Supporting Information**

**SI Methods**

**Participants.**

Participants of the current analyses were selected from the BOLD and READ projects. The BOLD project aimed to examine the neural trajectories underlying reading development in children with and without a family history of developmental dyslexia, while the goal of the READ project was to investigate whether the brain characteristics collected at the beginning of the kindergarten year might predict long-term reading achievements in children at behavioral risk of dyslexia. Therefore, the background information for assessing the familial risk of dyslexia was documented thoroughly in the BOLD project. For each family, we have collected comprehensive information on the dyslexia diagnoses, including which specific family members (mom, dad and/or siblings) have/had formal diagnoses, the date they were diagnosed, the location of the diagnoses, and their follow-up treatment plans (see similar protocol in previous longitudinal studies of dyslexia, e.g., Lyytinene et al., 2004; Scarborough, 1990; van der Leij et al., 2013). Based on this information, 39 FHD+ and 41 FHD- children were selected from the BOLD project. Moreover, to reach maximal sample size, an additional group of 13 children with family members formally diagnosed with dyslexia were further selected from the READ project.

**References.**

Lyytinen, H., Aro, M., Eklund, K., Erskine, J., Guttorm, T., Laakso, M. L., ... & Torppa, M. (2004). The development of children at familial risk for dyslexia: Birth to early school age. *Annals of dyslexia*, 54(2), 184-220.

Scarborough, H. S. (1990). Very early language deficits in dyslexic children. *Child development*, 61(6), 1728-1743.

van der Leij, A., Van Bergen, E., van Zuijen, T., De Jong, P., Maurits, N., & Maassen, B. (2013). Precursors of developmental dyslexia: an overview of the longitudinal Dutch dyslexia programme study. *Dyslexia*, 19(4), 191-213.

Figure S1

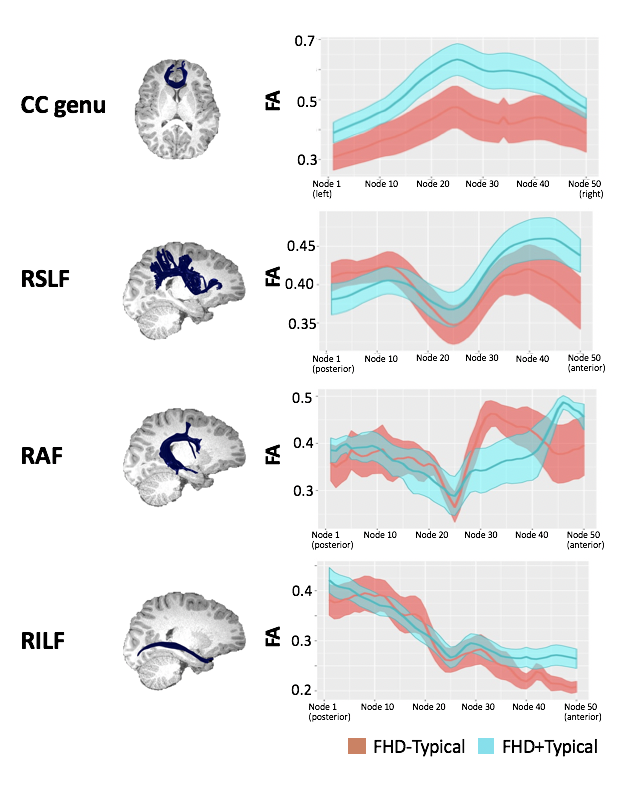

Figure S1. Tract profiles (FA values at all 50 nodes) in the corpus callosum genu and the three examined right-hemispheric tracts for FHD-Typical and FHD+Typical children. No significant group differences were observed in these tracts.

CC genu: the genu part of the corpus callosum; RSLF: superior longitudinal fasciculus; RAF: right arcuate fasciculus; RILF: right inferior longitudinal fasciculus.

FHD-Typical: children without family history of dyslexia who subsequently developed typical reading abilities; FHD+Typical: children with family history of dyslexia who subsequently developed typical reading abilities.

Table S1. Home literacy environment for FHD-TYP and FHD+TYP children.

| Environment | Options | FHD- Typical (%) | FHD+Typical (%) | Group effect  (Chi-square) | Significance |
| --- | --- | --- | --- | --- | --- |
| 1. Total number of parents/adult books in the home | 0-50 | 23.53 | 17.14 | 1.4 | 0.49 |
|  | 50-100 | 17.65 | 28.57 |  |  |
|  | 100+ | 50 | 42.86 |  |  |
|  | N/A | 8.82 | 11.43 |  |  |
| 2. Total number of children’s books in the home | 0-50 | 2.94 | 11.43 | 2.7 | 0.26 |
|  | 50-100 | 17.65 | 25.71 |  |  |
|  | 100+ | 67.65 | 54.29 |  |  |
|  | N/A | 11.76 | 8.57 |  |  |
| 3. Age (in months) of child when first read to (mean in months) | Prenatal | 2.94 | 2.86 | 6.1 | 0.52 |
|  | 0-3 | 58.82 | 60 |  |  |
|  | 3.1-6 | 14.71 | 11.43 |  |  |
|  | 6.1-9 | 0 | 0 |  |  |
|  | 9.1-12 | 0 | 2.86 |  |  |
|  | 12.1-24 | 0 | 2.86 |  |  |
|  | 24.1-36 | 0 | 0 |  |  |
|  | 36.1-48 | 0 | 2.86 |  |  |
|  | 48.1-60 | 5.88 | 0 |  |  |
|  | 60.1+ | 2.94 | 0 |  |  |
|  | N/A | 14.71 | 17.14 |  |  |
| 4. Amount of time at home that someone reads to child (hours/week) |  | 4.32 | 4.18 | 0.21¶ | 0.83 |
| 5. How often do family members read books, magazines or newspapers with the child? (times/week) | 1-2 | 0 | 0 | 7.3 | 0.03 |
|  | 3-4 | 8.82 | 8.57 |  |  |
|  | 5-6 | 8.82 | 34.26 |  |  |
|  | Daily | 76.47 | 48.57 |  |  |
|  | N/A | 5.88 | 8.57 |  |  |
| 6. How often do family members teach the child how to write? (times/week) | 1-2 | 29.41 | 25.71 | 0.17 | 0.98 |
|  | 3-4 | 32.35 | 34.29 |  |  |
|  | 5-6 | 14.71 | 17.14 |  |  |
|  | Daily | 14.71 | 14.29 |  |  |
|  | N/A | 8.82 | 8.57 |  |  |
| 7. How often do family members teach the child to count? (times/week) | 1-2 | 8.82 | 17.14 | 3.9 | 0.28 |
|  | 3-4 | 35.29 | 22.85 |  |  |
|  | 5-6 | 11.76 | 25.71 |  |  |
|  | Daily | 32.36 | 25.71 |  |  |
|  | N/A | 11.76 | 8.57 |  |  |
| 8. How often do family members help the child with their school work? | 1-2 | 8.82 | 17.14 | 4.8 | 0.19 |
|  | 3-4 | 20.59 | 11.43 |  |  |
|  | 5-6 | 8.82 | 25.71 |  |  |
|  | Daily | 23.53 | 17.14 |  |  |
|  | N/A | 38.34 | 28.57 |  |  |
| 9. How often do family members teach the child to read words? (times/week) | 1-2 | 26.47 | 22.86 | 1.4 | 0.71 |
|  | 3-4 | 11.76 | 17.14 |  |  |
|  | 5-6 | 11.76 | 8.57 |  |  |
|  | Daily | 8.82 | 17.14 |  |  |
|  | N/A | 41.18 | 34.29 |  |  |
| 10. How often does the child ask someone to read to them? (times/week) | 1-2 | 5.88 | 2.86 | 2.2 | 0.54 |
|  | 3-4 | 8.82 | 20 |  |  |
|  | 5-6 | 17.65 | 17.14 |  |  |
|  | Daily | 61.76 | 51.43 |  |  |
|  | N/A | 5.88 | 8.57 |  |  |
| 11. How often does someone at home help the child with their homework in reading and writing? (times/week) | 1-2 | 11.76 | 14.29 | 3.6 | 0.31 |
|  | 3-4 | 23.53 | 11.43 |  |  |
|  | 5-6 | 8.82 | 22.86 |  |  |
|  | Daily | 17.65 | 20 |  |  |
|  | N/A | 38.23 | 31.43 |  |  |
| 12. How often does the child look at books at home by themselves? (times/week) | 1-2 | 11.76 | 11.43 | 1.5 | 0.68 |
|  | 3-4 | 11.76 | 14.29 |  |  |
|  | 5-6 | 8.82 | 17.14 |  |  |
|  | Daily | 61.76 | 48.57 |  |  |
|  | N/A | 5.88 | 8.57 |  |  |
| 13. How often do family members read newspapers, books, or magazines? (times/week) | 1-2 | 5.88 | 14.29 | 1.8 | 0.62 |
|  | 3-4 | 5.88 | 5.71 |  |  |
|  | 5-6 | 11.76 | 14.29 |  |  |
|  | Daily | 70.59 | 57.14 |  |  |
|  | N/A | 5.88 | 8.57 |  |  |
| 14. How often do family members write messages, notes, or lists? (times/week) | 1-2 | 5.88 | 5.71 | 5.0 | 0.17 |
|  | 3-4 | 14.71 | 2.86 |  |  |
|  | 5-6 | 14.71 | 5.71 |  |  |
|  | Daily | 58.82 | 77.14 |  |  |
|  | N/A | 5.88 | 8.57 |  |  |
| 15. How often do family members write letters, cards, diaries, stories, or poems? (times/week) | 1-2 | 67.65 | 62.86 | 0.32 | 0.96 |
|  | 3-4 | 5.88 | 5.71 |  |  |
|  | 5-6 | 5.88 | 8.57 |  |  |
|  | Daily | 11.76 | 14.29 |  |  |
|  | N/A | 8.82 | 8.57 |  |  |
| 16. How often do family members share rhymes or jokes orally with the child? (times/week) | 1-2 | 26.47 | 14.29 | 2.6 | 0.45 |
|  | 3-4 | 32.36 | 25.71 |  |  |
|  | 5-6 | 11.76 | 20 |  |  |
|  | Daily | 23.53 | 31.43 |  |  |
|  | N/A | 5.88 | 8.57 |  |  |

For items with multiple choices, response frequency for each option (in percentage) was listed. Group effects were examined after excluding participants with no responses (i.e., Don’t know or N/A). Chi-square tests were performed on all items except question 4 (marked with ¶), where a two-sample t-test was applied.

FHD-Typical: children without family history of dyslexia who subsequently developed typical reading abilities; FHD+Typical: children with family history of dyslexia who subsequently developed typical reading abilities.

Table S2. Socioeconomic characteristics for FHD-TYP and FHD+TYP children.

| Characteristic | Option | FHD-Typical (%) | FHD+Typical (%) | Group effect  (Chi-square) | Significance |
| --- | --- | --- | --- | --- | --- |
| Mother characteristics | | | |  | |
| Education | 8th Grade or Less | 0 | 0 | 4.5 | 0.34 |
|  | Some High School | 0 | 0 |  |  |
|  | HS/GED | 2.94 | 11.43 |  |  |
|  | Associate’s Degree | 2.94 | 0 |  |  |
|  | Bachelor’s Degree | 35.29 | 40 |  |  |
|  | Master’s Degree | 41.18 | 31.43 |  |  |
|  | Doctorate or equivalent | 14.71 | 5.71 |  |  |
|  | N/A | 2.94 | 11.43 |  |  |
| Current Activity | Working full time | 32.35 | 20.00 | 0.49 | 0.78 |
|  | Working part time | 17.65 | 22.86 |  |  |
|  | Unemployed or laid off | 0 | 0 |  |  |
|  | Looking for work | 0 | 0 |  |  |
|  | Staying at home, raising a child | 47.06 | 45.71 |  |  |
|  | Retired | 0 | 0 |  |  |
|  | N/A | 2.94 | 11.43 |  |  |
| Money earned within the last 12 months | Less than $5,000 | 17.65 | 11.43 | 5.9 | 0.55 |
|  | $5,000-$11,999 | 5.88 | 0 |  |  |
|  | $12,000-$15,999 | 2.94 | 0 |  |  |
|  | $16,000-$24,999 | 0 | 0 |  |  |
|  | $25,000-$34,999 | 0 | 2.86 |  |  |
|  | $35,000-$49,000 | 8.82 | 2.86 |  |  |
|  | $50,000-$74,999 | 14.71 | 20.00 |  |  |
|  | $75,000-$99,999 | 14.71 | 8.57 |  |  |
|  | $100,000 and greater | 17.65 | 20.00 |  |  |
|  | Don’t know | 0 | 0 |  |  |
|  | N/A | 17.65 | 34.29 |  |  |
| Father characteristics | | | |  | |
| Education | 8th grade or Less | 0 | 0 | 5.8 | 0.22 |
|  | Some high school | 0 | 0 |  |  |
|  | HS/GED | 11.76 | 25.71 |  |  |
|  | Associate’s Degree | 5.88 | 11.43 |  |  |
|  | Bachelor’s Degree | 38.24 | 28.57 |  |  |
|  | Master’s Degree | 20.59 | 17.14 |  |  |
|  | Doctorate or equivalent | 20.59 | 5.71 |  |  |
|  | N/A | 2.94 | 11.43 |  |  |
| Current Activity | Working full time | 61.76 | 57.14 | 1.0 | 0.60 |
|  | Working part time | 2.94 | 2.86 |  |  |
|  | Unemployed or laid off | 0 | 0 |  |  |
|  | Looking for work | 0 | 2.86 |  |  |
|  | Staying at home, raising a child | 0 | 0 |  |  |
|  | Retired | 0 | 0 |  |  |
|  | No response | 35.29 | 37.14 |  |  |
| Family characteristics | | | |  | |
| Money earned within the last 12 months | Less than $5,000 | 0 | 0 | 4.2 | 0.24 |
|  | $5,000-$11,999 | 2.94 | 0 |  |  |
|  | $12,000-$15,999 | 0 | 0 |  |  |
|  | $16,000-$24,999 | 0 | 0 |  |  |
|  | $25,000-$34,999 | 0 | 0 |  |  |
|  | $35,000-$49,000 | 5.88 | 3.13 |  |  |
|  | $50,000-$74,999 | 0 | 0 |  |  |
|  | $75,000-$99,999 | 8.82 | 21.88 |  |  |
|  | $100,000 and greater | 14.71 | 6.25 |  |  |
|  | Don’t know | 55.88 | 46.88 |  |  |
|  | N/A | 11.76 | 21.88 |  |  |
| Length of time you could maintain standard of living if all income is lost | Less than 1 month | 11.76 | 2.86 | 3.6 | 0.47 |
|  | 1-2 months | 8.82 | 8.57 |  |  |
|  | 3-6 months | 29.41 | 25.71 |  |  |
|  | 7-12 months | 20.59 | 8.57 |  |  |
|  | More than 1 year | 20.59 | 28.57 |  |  |
|  | N/A | 8.82 | 25.71 |  |  |
| Home Owner Status | Occupied without payment of money/rent | 0 | 2.86 | 1.3 | 0.53 |
|  | Home rented for money | 11.76 | 8.57 |  |  |
|  | Home owned by you or being bought by household member | 82.35 | 62.86 |  |  |
|  | N/A | 5.88 | 25.71 |  |  |

For items with multiple choices, response frequency for each option (in percentage) was listed. Chi-square tests were performance to evaluate the group differences after excluding participants with no responses (i.e., Don’t know or N/A).

FHD-Typical: children without family history of dyslexia who subsequently developed typical reading abilities; FHD+Typical: children with family history of dyslexia who subsequently developed typical reading abilities.

Table S3. Averaged reading performance across scores available for all the grades.

|  | FHD-Typical | FHD+Typical | Group Effect |
| --- | --- | --- | --- |
| WRMT-R: Word ID | 112 ± 9.4 | 109 ± 11 | *t*_67_ = 1.1; *p* = 0.3 |
| WRMT-R: Word Attack | 110 ± 9.9 | 111 ± 10 | *t*_67_ = 0.08; *p* = 0.9 |
| TOWRE: Sight Word Efficiency | 109 ± 12 | 104 ± 11 | *t*_67_ = 1.8; *p* = 0.08 |
| TOWRE: Phonemic Decoding Efficiency | 104 ± 11 | 103 ± 9.3 | *t*_67_ = 0.4; *p* = 0.7 |

Note. Standard scores were reported for all the assessments. No significant differences were observed between the FHD-Typical and FHD+Typical children.

FHD-Typical: children without family history of dyslexia who subsequently developed typical reading abilities; FHD+Typical: children with family history of dyslexia who subsequently developed typical reading abilities; WRMT-R: Woodcock Reading Mastery Tests-Revised; TOWRE: Test of Word Reading Efficiency.

Table S4. FMRI experiment performance conducted at the pre-reading stage

|  | **FHD-Typical** | **FHD+Typical** | **Group Effect** |
| --- | --- | --- | --- |
| Head Motion: Left to right (mm) | 0.045 ± 0.37 | 0.065 ± 0.34 | *t*_58_ = 0.2; *p* = 0.8 |
| Head Motion: Posterior to anterior (mm) | 0.29 ± 0.81 | -0.048 ± 0.77 | *t*_58_ = 1.6; *p* = 0.1 |
| Head Motion: Bottom to top (mm) | 0.44 ± 1.5 | 0.63 ± 1.8 | *t*_58_ = 0.4; *p* = 0.7 |
| Head Motion: Pitch (°) | 0.17 ± 1.7 | 0.33 ± 2.0 | *t*_58_ = 0.3; *p* = 0.7 |
| Head Motion: Yaw (°) | -0.21 ± 0.41 | -0.13 ± 0.55 | *t*_58_ = 0.7; *p* = 0.5 |
| Head Motion: Roll (°) | 0.12 ± 0.73 | -0.18 ± 0.44 | *t*_58_ = 2.0; *p* = 0.06 |
| # of correct responses | 17 ± 7.2 | 17 ± 6.3 | *t_54_* = 1.4; *p* = 0.2 |
| Response times (seconds) | 2336 ± 480 | 2170 ± 422 | *t*_54_ = 0.1; *p* = 0.9 |

Note. No significant group effect was observed for head movement and task performance during the fMRI experiment.

Video. 3D render of the examined white matter tracts displayed for one representative participant.

Green: right superior longitudinal fasciculus; Blue: right acute fasciculus; yellow: right inferior longitudinal fasciculus; purple: corpus callosum
